# Simulated field potentials capture multiple simultaneous processes underlying human temporal attention

**DOI:** 10.64898/2026.09.17.752423

**Authors:** Sydney E. Smith, Dillan Cellier, Alison Rigby, Burke Q. Rosen, Jacob Garrett, Alexander J. Simon, Sharona Ben-Haim, Jerry J. Shih, Keith B. Doelling, Bradley Voytek

**Author notes:** These authors jointly supervised the work (Keith B. Doelling, Bradley Voytek).

## Abstract

Essential cognitive processes like attention, memory, and decision-making require the brain to perform many operations simultaneously. Disentangling the neural mechanisms underlying these processes is challenging because they often unfold concurrently with similar dynamics. In auditory temporal expectation, these mechanisms include the accumulation of incoming sensory information in the auditory cortex interacting with cortical signals biasing activity in anticipation of future stimuli. Here, we leverage invasive human field potential recordings from participants performing a subjective temporal expectation task. Rather than relying on traditional neuroimaging methods that average or transform neural responses, we instead simulate each trial’s predicted field potential, based on known field potential physiology combined with components of established cognitive theories of the temporal expectation task: sensory input, top-down bias, and sensory anticipation. Our predicted field potentials strongly correlate with neural activity in auditory regions and in regions associated with auditory temporal perception, including the parietal, temporal, and prefrontal cortices. By differentiating individual cognitive mechanisms through simulation, we identify neural signatures of attentional bias and anticipatory sensory activity across multiple cortical regions that predict trial-by-trial perceptual accuracy and confidence.

## Main

Adaptive behavior depends on the coordinated interaction of multiple cognitive processes that unfold simultaneously, and this is critical for our daily function. As you read the words on this page, sensory input from your retina is rapidly processed through your visual and language systems, allowing you to predict what the missing word is at the end of this []^1^. Rather than operating independently, these cognitive processes interact continuously to shape perception and behavior^2^. While adaptive, the co-occurrence of these cognitive functions in time makes studying their neural underpinnings in humans challenging. Neural activity recorded during behavior often reflects the combined influence of many concurrent computations distributed across overlapping brain networks^3,4^. This overlap presents a fundamental challenge for cognitive neuroscience: determining which aspects of neural activity correspond to which cognitive processes. Even in paradigms designed to isolate specific functions, like studies of rhythm perception and auditory temporal expectation^5^, multiple processes must co-occur.

Several simultaneous neurocognitive mechanisms likely support temporal predictions in auditory perception. One component of this process is the bottom-up sensation of sound itself, where stimulus-selective cortices respond to incoming auditory afferents from sensory pathways^6,7^. Concurrently, as a listener gains more experience with temporal regularities in the environment, their expectations can drive selective attention in a top-down fashion, modulating neural activity in anticipation of upcoming stimuli^8^. Crucially, attentional processes must integrate sensory inputs in neural circuits to update predictions and guide auditory timing behavior.

Efforts to understand the behavioral and neural mechanisms by which inputs from external sources can be dynamically integrated with internal attentional processes have produced a burgeoning field over the past several decades^5,9–13^. While the neural correlates of temporal expectation have been studied in using different types of timing structures – hazards, cued associations, and rhythms, to name a few – nearly all have identified a mechanistic role of neural excitability modulation, where prior to an anticipated stimulus neurons in the relevant sensory cortex are more likely to spike^8^. In mice, rats, and non-human primates, firing rates in sensory cortices increase in anticipation of upcoming stimuli^14–17^. Evidence from non-invasive neuroimaging in humans also supports neural engagement in selective attentional processes via dynamic modulations of neural oscillations^5,18,19^ in sensory cortices as well as prefrontal and parietal regions^20^. Accompanying these experimental advances, a wealth of computational models have been developed to hypothesize how temporal prediction operates for different temporal structures^21^ and timescales^22^, largely validated on behavioral findings^13,23–26^.

A persistent challenge in studying temporal expectation in humans is the inability to isolate sensory responses from the attentional excitability modulations in measurements of neural activity that represent the aggregation of countless physiological processes^27^. How do we tell top-down apart from bottom-up? How are they integrated together in neural circuitry? How do we detect each process in neural signals? And finally, how does each component contribute to temporal expectation behavior?

Answering these questions is challenging, as the processes of bottom-up sensation, top-down attentional bias, and their integration in neural circuits are all simultaneous, interdependent, and appear highly similar in neural signals^28,29^. In some cases, approaches like trial averaging and bandpass filtering, which are intended to make neural signals more interpretable, can unintentionally obscure or distort natural temporal variability of cognitive processes^30^ and the neural signals themselves, especially when they violate assumptions of periodicity and sinusoidality^31,32^. Furthermore, with non-invasive neuroimaging like electroencephalography (EEG) and functional magnetic resonance imaging (fMRI), results are largely constrained to informing us of what brain regions are associated with temporal prediction (using fMRI), or when the prediction of a temporal event occurs (using EEG), but are rarely able to sufficiently model both.

Here, we present a new experimental and analytical approach to address these issues: a simulation-based method that combines cognitive modeling with field potential physiology. With this approach, we differentiate multiple concurrent mechanisms that contribute to temporal expectation in invasive cortical field potential recordings from ten human participants performing a temporal prediction task. In this task, participants listened to a sequence of three to five tones presented isochronously at 1-3 Hz. The last tone in each trial was temporally jittered relative to the isochronous tone timing, and participants were tasked with characterizing it as having occurred earlier or later than expected, compared to the isochronous pattern. Perceptual accuracy of the target tone improved with more prior tones and longer jitters (**Fig. 1**).

**Figure 1.**
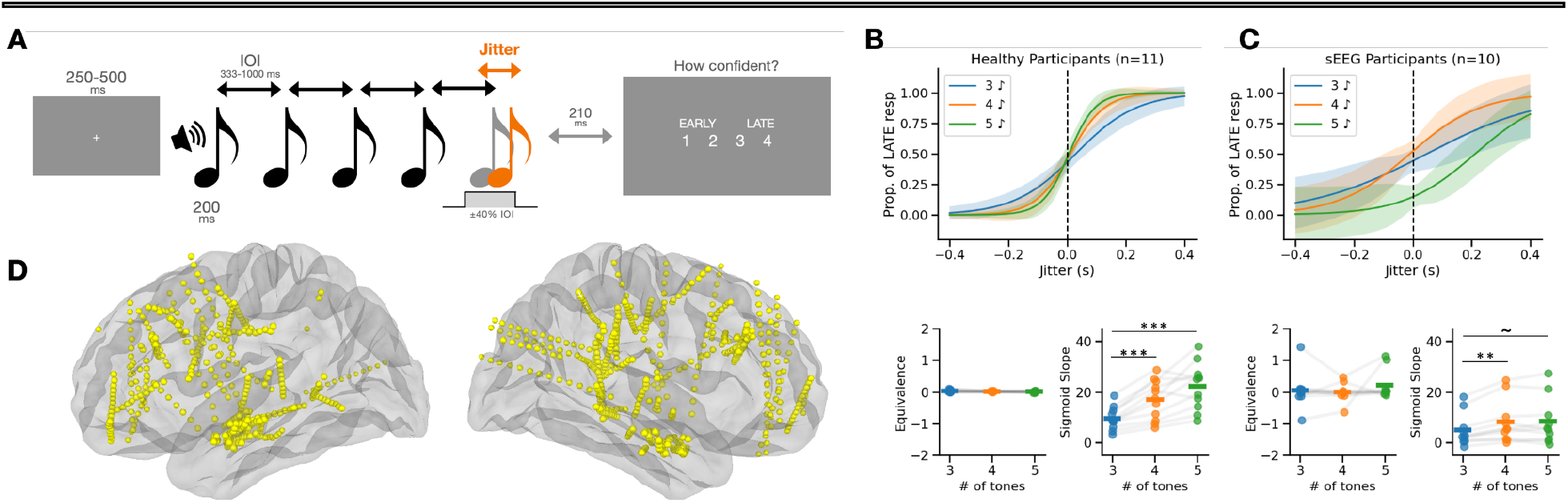
Task, behavior, and electrode locations. **(A)** Task design. A fixation period of 250-500 ms is followed by the presentation of 3-5 piano tones (C4 - B4). The first 2-4 tones are presented isochronously with inter-onset intervals ranging from 333-1000 ms (1-3 Hz). The last tone in the sequence is jittered relative to the inter-onset interval (IOI). Trial-wise jitters were randomly drawn from a uniform distribution ranging ±40% of the IOI. After 210 ms, the participant is cued to respond if the tone is early or late, and how confident they are in their response on a scale of 1-4, where a confident early response is indicated by a keypress of 1 and a confident late response is indicated by keypress of 4. A less confident response is a forced choice between 2 and 3. **(B)** Average sigmoid fits of behavioral accuracy in healthy participants performing the task, by condition (blue: 3 tones, orange: 4 tones, green: 5 tones). Average fit is a solid line and shading represents one standard deviation. Equivalence point (*μ*) of sigmoid fits did not differ between conditions (one-way rm-ANOVA), but slope (*σ*) significantly increased on trials with more than three tones (one-way rm-ANOVA: p = 2.30×10^-5^; *σ*_3-tone_ = 9.56±4.23, *σ*_4-tone_ = 16.90±7.46, *σ*_5-tone_ = 22.23±8.71; 3 vs. 4: p_corr_ = 6.50×10^-4^; 3 vs. 5: p_corr_ = 6.09×10^-4^; n = 11). **(C)** Average sigmoid fits of behavioral accuracy in sEEG participants performing the task. Color, line style, and shading like panel B. *μ* did not differ between conditions (one-way rm-ANOVA). *σ* was significantly greater on three-tone trials than four-tone trials with trending differences between three-and five-tone trials (one-way rm-ANOVA: p = 3.33×10^-3^; *σ*_3-tone_ = 4.95±6.10, *σ*_4-tone_ = 8.24±8.15, *σ*_5-tone_ = 8.26±8.75; 3 vs. 4: p_corr_ = 8.62×10^-3^; 3 vs. 5: p_corr_ = 0.054; n = 10). *σ* was significantly greater in healthy participants than sEEG participants (one-way mixed ANOVA: p = 2.36 x 10^-4^). **(D)** Cortical locations of midpoints between bipolar contact pairs in ten sEEG participants, where electrodes were implanted based solely on clinical needs.

Instead of transforming the neural activity with filters or averaging neural responses across trials, we simulated neural spike trains unique to each individual trial’s characteristics, preserving each trial’s timescale (**Fig. 2A**). Spike trains were simulated using three different models: (1) a sensory input model derived from auditory pathway physiology that results in a rapid increase in firing rate at tone onset that decays, but sustains at above-baseline levels, throughout the duration of the tone^7^; (2) a top-down bias model that reflects accumulating certainty of the timing of auditory events^33^ and; (3) an anticipation model that simulates excitation-inhibition dynamics to anticipate future events^25^. Because our data do not include single neuron spiking activity, but rather aggregate local field potential (LFP) recordings, we simulated each trial’s predicted LFP, for each model, based on the known physiology of the LFP (**Fig. 2B**). Specifically, the LFP largely reflects the aggregate transmembrane currents near the electrode^27^, which allows us to predict each trial’s LFP from the convolution of an excitatory postsynaptic current with our simulated spiking models^34^. While we have not tested our approach against the entirety of the rich collection of models proposed in temporal prediction (including oscillatory or timekeeper models), by developing this approach, we demonstrate the ability to directly relate underlying models of temporal prediction to the physiological data they might support.

**Figure 2.**
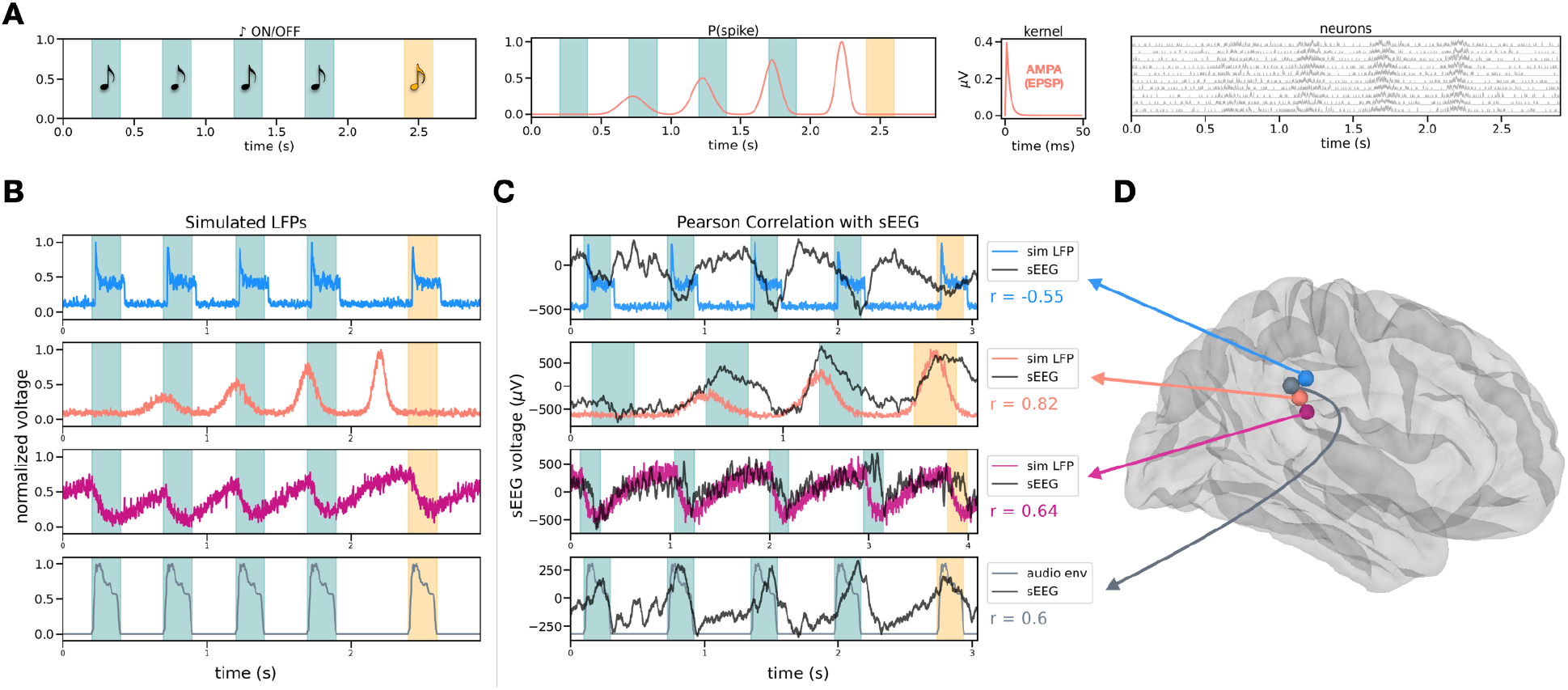
Trial-by-trial spiking-to-LFP simulation and correlation with sEEG. **(A)** Trial-wise LFP simulation technique. LFPs are simulated to represent each model under the conditions of a given trial. The onsets of tones in each trial are determined by the trial parameters (number of tones, IOI, and jitter). The trial depicted here has five tones, an IOI of 500 ms, and a target tone jitter of 200 ms. Each tone is 200 ms long. The probability of a spike occurring during the trial is generated for each model given the trial parameters (see Methods). The spike probability signal is then interpolated with a population of 100 randomly-spiking neurons (ten are shown here). Each spike is convolved with a voltage kernel representing the excitatory postsynaptic potential of an AMPA receptor, reflecting both corticopetal excitatory drive, and top-down excitatory drive from association cortex, onto auditory cortex. The convolved spiking signals for all neurons are then summed to generate the simulated LFP for the trial (see panel B). **(B)** Simulated LFPs for the trial depicted in panel A for each model: Bottom-up sensory input (blue), top-down bias (salmon), and the SAM_y_ dynamical systems model (magenta). The auditory envelope (control) of the same trial is shown in gray. **(C)** Comparison of simulated LFPs with sEEG. Four trials from four different electrodes are depicted, each with the simulated LFP (colors same as panel B) and the raw, unfiltered sEEG signal (black). Voltages from simulations are normalized between zero and one, like panel B. sEEG voltages are displayed in microvolts, as recorded from the bipolar contact pair midpoint. The Pearson correlation coefficient (*r*) represents the strength of the similarity between the two signals, where a strong negative correlation indicates a signal with opposite voltage, suggesting a putative dipole (see blue, bottom-up signal). These trials were selected for visualization for their high similarity values, and are not representative of the average electrode. **(D)** Estimated cortical location of sEEG depth electrodes, the sources of the sEEG signals (black) depicted in panel C. Colors indicate the source of the sEEG signals compared to the simulated LFPs displayed in panel C.

Remarkably, the predicted LFP signal for each trial strongly correlated with the actual LFP across the cortical auditory perceptual and temporal prediction network, including superior temporal, prefrontal, and parietal cortices (**Fig. 2C**). These correlations significantly exceeded a control correlation with the amplitude envelope of the presented audio signal for two out of three models tested. While both top-down and sensory anticipation models were most strongly related to prefrontal cortical activity, the sensory anticipation model better explained the data in the superior temporal cortex (**Fig. 3**). We also show that, when the correlation between cortical LFP signal and the sensory anticipation model was stronger, behavioral accuracy was better (**Fig. 4**). This kind of observation is markedly different from traditional human neuroscience approaches where a signal average correlates with behavior. Instead of examining the fMRI BOLD signal, EEG event related potentials or oscillatory band power, or even intracranial high frequency power, we instead show that the more closely a given trial’s raw LFP corresponds to the sensory anticipation model of temporal expectation, the more accurate the person’s assessment of whether the target tone was earlier or later than expected.

**Figure 3.**
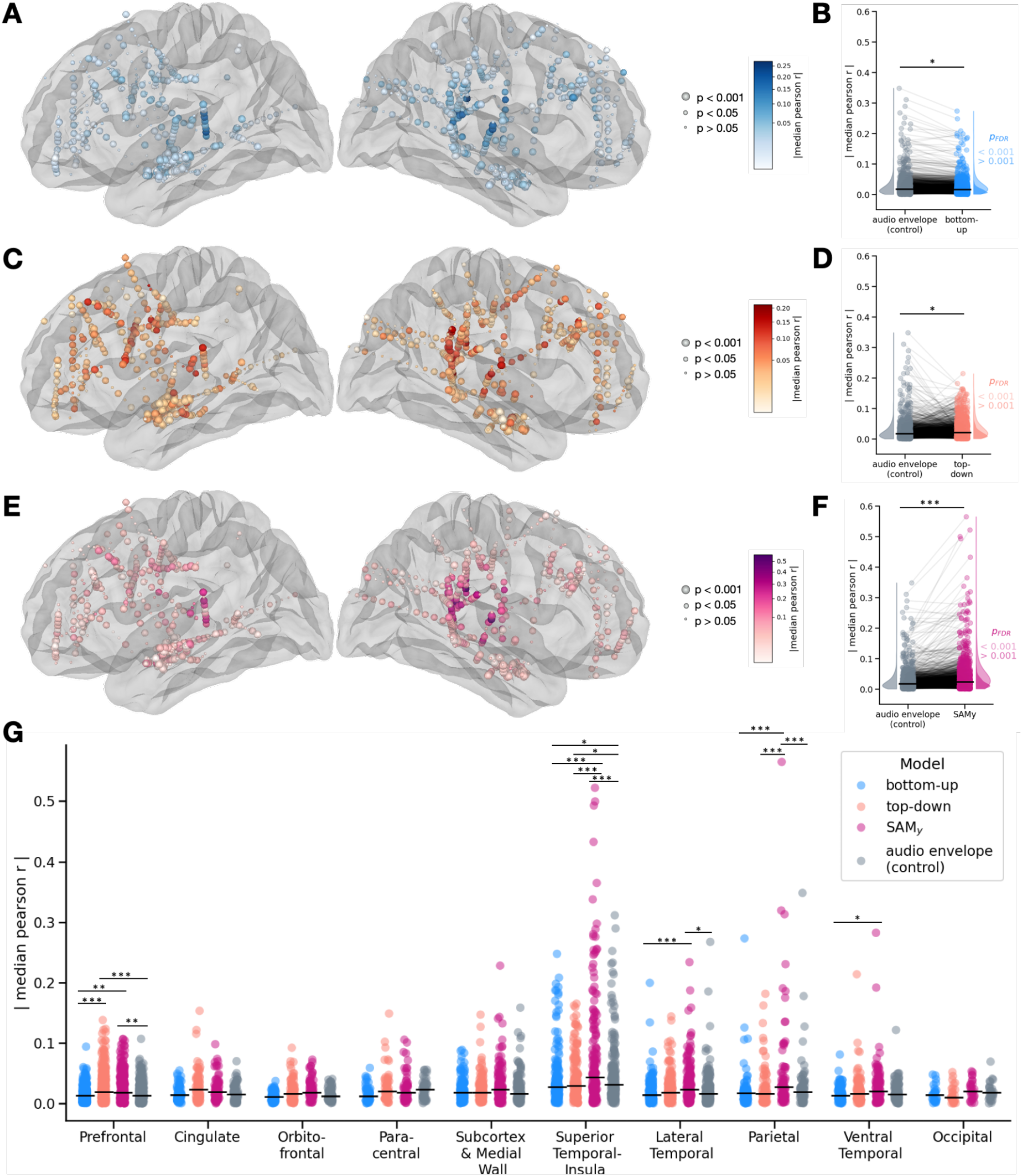
Pearson correlation strength of sEEG to simulated LFP. **(A)** Bottom-up sensory input model similarity to neural signals. sEEG electrode color intensity represents the similarity of the neural signal to the LFP simulated from the bottom-up model. Similarity is quantified as the absolute value of the median Pearson coefficient (*r*) across all included trials. Electrode size represents the FDR-corrected p-value from a 1000-fold permutation test. **(B)** In a linear mixed-effects model with participant as a random effect, there was a significant main effect of model type on similarity across all cortical electrodes (n = 1258, omnibus Wald test: *χ*^2^(3) = 138.3, p < 10^-5^). The bottom-up model was significantly less similar to sEEG than the audio envelope (r_bottom-up_ EMM = 0.021; r_control_ = 0.025; pairwise Wald test: p_FDR_ = 0.02). **(C)** Top-down bias model similarity to neural signals. **(D)** The top-down model was significantly more similar to sEEG than the audio envelope (r_top-down_ = 0.028; r_control_ = 0.026; pairwise Wald test test: p_FDR_ = 0.032) and the bottom-up model (r_top-down_ = 0.028; r_bottom-up_ = 0.021; pairwise Wald test: p_FDR_ < 10^-5^. **(E)** Dynamical systems SAM_y_ (y-unit) model similarity to neural signals. **(F)** SAM_y_ had significantly higher similarity values than the audio envelope control condition (r_SAMy_ = 0.037; r_control_ = 0.025; pairwise Wald test: p_FDR_ < 10^-5^). SAM_y_ had higher similarity values than both the top-down model (r_SAMy_ = 0.037; r_top-down_ = 0.028; pairwise Wald test test: p_FDR_ < 10^-5^) and the bottom-up model (r_SAMy_= 0.037; r_bottom-up_ = 0.021; pairwise Wald test: p_FDR_ < 10^-5^). **(G)** Similarity values of all electrodes by cortical region. Absolute value of the median *r* coefficient is plotted by cortical region, based on a coarse regional atlas. Horizontal black bars represent region-wise medians. Estimated marginal means for all regions and Wald-z tests for within-region model comparisons are available in **Table 3 and Supp. Table 1**. See **Supp. Fig. 3** for paired plots of model and control similarity values in significant regions. All p-values are FDR-adjusted for multiple comparisons. Stars indicate significant differences between models within a region (∗ p < 0.05, ∗∗ p < 0.01, ∗∗∗ p < 0.001). All participants had sEEG coverage in the prefrontal and superior temporal cortex, but sEEG from other regions represents a subset of participants.

**Figure 4.**
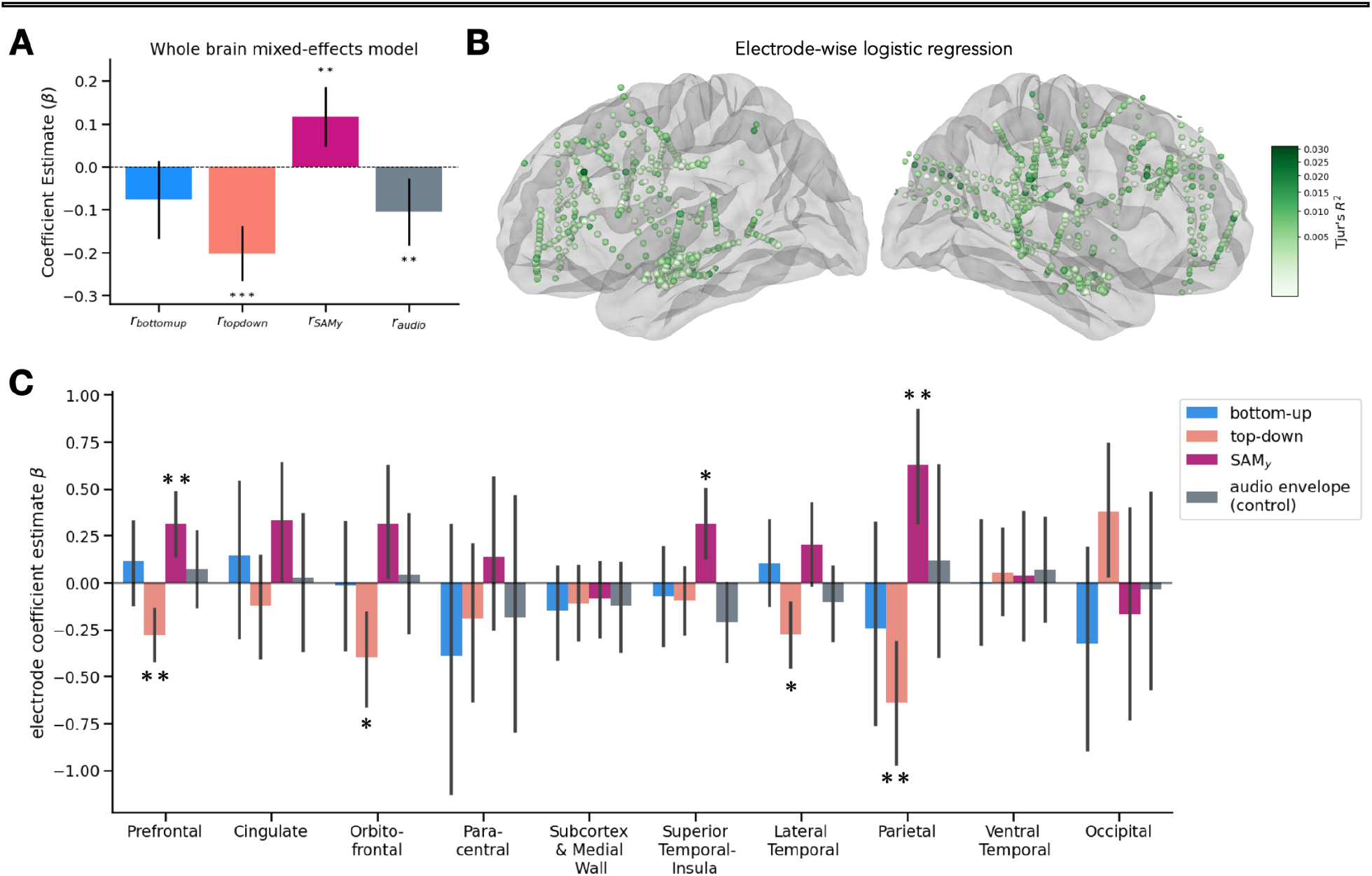
Mixed-effects modeling of trial-wise accuracy from neural model similarity. **(A)** Estimated slopes (β) relating neural fit (|r|) to behavioral accuracy for each model type, derived from a mixed-effects logistic regression with participant as a random intercept (Tjur’s R^2^ = 0.06). Bars indicate fixed-effect slopes from estimated marginal means; error bars represent 95% CI. The slope for SAM_y_ model was significantly positive (β_SAMy_ = 0.12, p = 1.05 x 10^-3^), while the top-down model (β_top-down_ =-0.20, p < 10^-5^) and audio envelope (β_audio_=-0.11, p = 0.01) were significantly negative. **(B)** Cortical map of electrode-wise logistic regression of accuracy in all participants. Color represents Tjur’s R^2^. **(C)** Estimated slopes (β) of electrode-wise regressions shown in B, distributed by cortical region. Bars indicate slopes from estimated marginal means; error bars represent 95% CI. Stars indicate significantly positive/negative slopes for electrodes in that region (one-sample t-test: ∗ p_FDR_ < 0.05, ∗∗ p_FDR_ < 0.01).

**Table 1.** Patient demographics and characteristics. ‘Yrs Music’ refers to the number of years of musical training and/or experience. Channels were excluded based on clinical indication for proximity to pathological tissue. Epochs were excluded if they contained interictal activity that spread beyond excluded channels.

| ID | Age | Sex | Hand | Yrs Music | sEEG chs | chs excluded | epochs excluded | Dx | Pathology |
| --- | --- | --- | --- | --- | --- | --- | --- | --- | --- |
| 1 | 30s | F | R | 1 | 142 | 9 | 39 | Lesional right frontal lobe epilepsy | Periventricular heterotopia |
| 2 | 20s | M | R | 1 | 168 | 33 | 12 | Lesional right temporal lobe epilepsy | Focal cortical dysplasia |
| 3 | 30s | M | R | 0 | 162 | 28 | 1 | Non-lesional post-traumatic bilateral temporal lobe epilepsy |  |
| 4 | 30s | M | L | 0 | 140 | 15 | 0 | Lesional post-traumatic bilateral temporal epilepsy | Post-traumatic encephalomalacia |
| 5 | 20s | M | R | 0 | 238 | 35 | 11 | Lesional right temporal lobe epilepsy | Right choroid fissure cyst, post-traumatic left frontal encephalomalacia |
| 6 | 20s | M | R | 15 | 194 | 16 | 16 | Lesional right temporal lobe epilepsy | Nodular heterotopia |
| 7 | 30s | F | R | 1 | 146 | 36 | 8 | Non-lesional left temporal lobe epilepsy |  |
| 8 | 40s | F | R | 0 | 174 | 21 | 0 | Lesional right temporal lobe epilepsy | Nodular heterotopia |
| 9 | 20s | F | R | 4 | 212 | 22 | 59 | Lesional right frontal lobe epilepsy | Nodular heterotopia |
| 10 | <20 | M | R | 7 | 234 | 59 | 35 | Non-lesional right temporal lobe epilepsy |  |

**Table 2.** Coarse cortical parcellation groupings. Cortical regions from the Desikan cortical atlas were grouped into coarse regions, a technique similar to ^35^Jiang et al. 2019.

| Combined ROI | Original Freesurfer Label (Desikan Atlas) | # participants | # electrodes |
| --- | --- | --- | --- |
| Prefrontal | pars opercularis, pars triangularis, caudal middle frontal, rostral middle frontal, superior frontal | 10 | 338 |
| Cingulate | caudal anterior cingulate, posterior cingulate, isthmus cingulate | 9 | 79 |
| Orbitofrontal | medial orbitofrontal, lateral orbitofrontal, frontal pole, pars orbitalis | 7 | 83 |
| Paracentral | postcentral, precentral, paracentral | 6 | 53 |
| Subcortex & Medial Wall | unknown | 9 | 136 |
| Superior Temporal-Insula | superior temporal, insula | 10 | 177 |
| Lateral Temporal | middle temporal, banks of superior temporal gyrus, inferior temporal | 9 | 178 |
| Parietal | inferior parietal, supramarginal, precuneus | 5 | 77 |
| Ventral Temporal | lingual, fusiform, parahippocampal | 8 | 110 |
| Occipital | lateral occipital, pericalcarine | 2 | 27 |

**Table 3.** Estimated marginal means of model sEEG similarity. . Whole brain EMMs are computed from a LME analysis of |median(Pearson *r*)| with model type as a fixed effect and participant as a random intercept. Region-wise EMMs are computed from an LME analysis of |median(Pearson *r*)| with the interaction of model type and region as a fixed effect and participant as a random intercept.

| region | # participants | # electrodes | model | r (EMM) | CI (2.5%) | CI (97.5%) |
| --- | --- | --- | --- | --- | --- | --- |
| Whole Brain | 10 | 1258 | top-down | 0.028 | 0.022 | 0.033 |
|  |  |  | bottom-up | 0.021 | 0.016 | 0.026 |
|  |  |  | SAM<sub>y</sub> | 0.037 | 0.032 | 0.042 |
|  |  |  | audio env | 0.025 | 0.019 | 0.03 |
| Prefrontal | 10 | 338 | top-down | 0.027 | 0.022 | 0.033 |
|  |  |  | bottom-up | 0.015 | 0.01 | 0.021 |
|  |  |  | SAM<sub>y</sub> | 0.025 | 0.019 | 0.03 |
|  |  |  | audio env | 0.017 | 0.011 | 0.022 |
| Cingulate | 9 | 79 | top-down | 0.03 | 0.021 | 0.038 |
|  |  |  | bottom-up | 0.017 | 0.009 | 0.026 |
|  |  |  | SAM<sub>y</sub> | 0.025 | 0.016 | 0.033 |
|  |  |  | audio env | 0.018 | 0.009 | 0.026 |
| Orbitofrontal | 7 | 83 | top-down | 0.022 | 0.014 | 0.031 |
|  |  |  | bottom-up | 0.013 | 0.004 | 0.021 |
|  |  |  | SAM<sub>y</sub> | 0.024 | 0.016 | 0.033 |
|  |  |  | audio env | 0.016 | 0.007 | 0.024 |
| Paracentral | 6 | 53 | top-down | 0.027 | 0.017 | 0.037 |
|  |  |  | bottom-up | 0.014 | 0.004 | 0.024 |
|  |  |  | SAM<sub>y</sub> | 0.026 | 0.016 | 0.036 |
|  |  |  | audio env | 0.021 | 0.011 | 0.031 |
| Parietal | 5 | 77 | top-down | 0.025 | 0.016 | 0.034 |
|  |  |  | bottom-up | 0.021 | 0.012 | 0.029 |
|  |  |  | SAM<sub>y</sub> | 0.051 | 0.043 | 0.06 |
|  |  |  | audio env | 0.025 | 0.017 | 0.034 |
| Medial Wall & Subcortex | 9 | 136 | top-down | 0.025 | 0.018 | 0.032 |
|  |  |  | bottom-up | 0.024 | 0.017 | 0.031 |
|  |  |  | SAM<sub>y</sub> | 0.032 | 0.025 | 0.039 |
|  |  |  | audio env | 0.026 | 0.019 | 0.033 |
| Superior Temporal-Insula | 10 | 177 | top-down | 0.040 | 0.034 | 0.046 |
|  |  |  | bottom-up | 0.042 | 0.035 | 0.048 |
|  |  |  | SAM<sub>y</sub> | 0.081 | 0.075 | 0.088 |
|  |  |  | audio env | 0.051 | 0.044 | 0.057 |
| Lateral Temporal | 9 | 178 | top-down | 0.026 | 0.019 | 0.032 |
|  |  |  | bottom-up | 0.020 | 0.014 | 0.027 |
|  |  |  | SAM<sub>y</sub> | 0.034 | 0.028 | 0.041 |
|  |  |  | audio env | 0.024 | 0.018 | 0.031 |
| Ventral Temporal | 8 | 110 | top-down | 0.023 | 0.016 | 0.031 |
|  |  |  | bottom-up | 0.017 | 0.009 | 0.024 |
|  |  |  | SAM<sub>y</sub> | 0.030 | 0.023 | 0.038 |
|  |  |  | audio env | 0.019 | 0.012 | 0.027 |
| Occipital | 2 | 27 | top-down | 0.017 | 0.004 | 0.031 |
|  |  |  | bottom-up | 0.018 | 0.004 | 0.031 |
|  |  |  | SAM<sub>y</sub> | 0.025 | 0.011 | 0.039 |
|  |  |  | audio env | 0.022 | 0.008 | 0.035 |

### Auditory temporal expectation task

Eleven healthy participants and ten patients with implanted intracranial electrodes performed an auditory temporal expectation task (**Fig. 1A**). In the task, participants were presented with a sequence of three to five piano tones. Tones were isochronous, with equal inter-onset intervals (IOIs) except for the last tone in the sequence, the target tone, which was jittered. At the end of each trial, participants were prompted to make a timing judgement about whether the target tone was early or late. We hypothesized that behavioral accuracy would improve with a greater number of isochronous tones, with lowest accuracy on three-tone trials, and highest accuracy on five-tone trials.

To model behavioral accuracy on the temporal expectation task, a sigmoid function was fit to a participant’s responses binned by jitter. The sigmoid function is parameterized by its slope (*σ*) and equivalence point (*μ*), where *σ* corresponds to overall behavioral accuracy and *μ* indicates a bias towards responding EARLY or LATE. In eleven healthy controls performing the temporal expectation task, timing judgement accuracy significantly improved with a greater number of prior tones (**Fig. 1B**), as quantified by *σ*. Specifically, *σ* (accuracy) increased between the three-and four-tone conditions and between the three-and five-tone conditions, with trending increases between the four-and five-tone conditions. sEEG participants’ behavior was more variable (**Fig. 1C)**; only the three-and four-tone conditions showed significant differences in *σ*. Slope was significantly greater in healthy controls compared to sEEG participants, and this effect was driven by three sEEG participants who performed near chance (**Supp. Fig. 1**).

### Simulation of predicted field potentials

We simulated LFPs based on three different computational models of the hypothesized neural processes underlying auditory temporal expectation (**Fig. 2**). These included a model of bottom-up sensory input, a model of top-down bias, and a dynamical systems model of sensory anticipation.

In the bottom-up sensory input model, excitatory drive directly reflects incoming stimuli, exponentially increasing at the onset then plateauing through the duration of each tone, as derived from physiological models of the auditory system^7^. In the top-down bias model, prefrontal excitatory drive fluctuates in advance of the auditory stimulus arriving in auditory cortical regions. This can be modeled as a sequence of narrowing, heightening Gaussians, with peaks aligned to each tone onset in the trial. Finally, in the dynamical systems sensory anticipation model (SAM_y_), excitatory drive adaptively ramps in anticipation of an expected stimulus, reaching a threshold at the expected onset of the incoming sensory stimulus. See **Methods** for a complete description of each model.

On any given trial of the temporal expectation task, the trial-specific features (number of tones, IOI, and jitter) were input into each of the three models to generate a probability signal consistent with that model’s conditions (**Fig. 2A**, left). This probability signal *P(spike)* represented the likelihood of an excitatory postsynaptic potential occurring at any given moment during the trial. For instance, in the top-down bias model, the probability of a spike occurring at the onset of a tone is high, and the probability of a spike occurring shortly after a tone is low. When this probability signal was interpolated with a randomly-firing neuron, spikes were distributed based on the events in the trial. Here, all spikes were treated as excitatory inputs into the neural population participating in the cognitive operation of the rhythmic attention task. Therefore, spikes were convolved with a kernel representing the postsynaptic potential of an AMPA receptor, the primary glutamate receptor of cortical neurons (**Fig. 2A**, right). Convolving the neuron’s spikes with the AMPA kernel produced a voltage trace for a single neuron. Summing the voltage traces of 100 spiking neurons returned a putative LFP. Note, however, that the exact timescale of the convolved kernel – within reasonable physiological bounds of 2 - 200 ms – does not have a significant impact on the subsequent neurobehavioral results, although the strength of the correlation between the sEEG signal and the top-down and bottom-up models improves with longer kernel timescales (see **Supplementary Methods**). The amplitude envelope of the audio stimulus on each trial was used as a control condition. Given that the exogenous stimulus exhibited similar rhythmicity to all three models, this control accounts for that, but without the assumption of an endogenous physiological process.

With this procedure, we simulated three putative LFPs for each trial on the task, each representing the hypothesized neural population activity described by the corresponding model (**Fig. 2B**). On any given trial, the three simulated LFPs, and the amplitude envelope of the auditory stimuli, were directly compared to the raw bipolar-referenced sEEG signal recorded on that trial from all included electrodes using a Pearson correlation (**Fig. 2C**). A coefficient of zero suggests no similarity and a coefficient of ±1 suggests a high degree of similarity between the signals. A negative correlation coefficient indicates a signal with similar temporal dynamics, but an opposite voltage from the simulated LFP, potentially representing a neural dipole in the sEEG (**Fig. 2C, top**). Figure 2C depicts four trials recorded from one participant with coverage in parietal, superior temporal, and insular cortical regions (**Fig. 2D**). The trials shown exhibited a high degree of similarity between the simulated LFP and the recorded sEEG signal and are not representative of the variability in model-sEEG similarity between electrodes (**Supp. Fig. 2**).

### Predicted field potentials correlate with sEEG in prefrontal, temporal, and parietal cortex

To investigate the spatial properties of observed similarity values, LFPs were compared to sEEG at every available depth electrode (**Fig. 1C**). For each included, artifact-free trial, an LFP was simulated for each model and compared with sEEG data using a Pearson correlation. To assign statistical significance to correlations, a 1000-fold permutation analysis was performed, where LFPs were simulated with randomly-shuffled trial parameters (number of tones, IOIs, and jitters), then correlated to the sEEG at each electrode. Permutation p-values were corrected for multiple comparisons. An sEEG signal’s similarity to a model was described by the absolute value of the median Pearson coefficient of all trials at that electrode. Absolute value was used to accommodate for electrophysiological dipoles, which are observed often in sEEG, and would generate a highly negative Pearson coefficient when a signal has opposite voltage but similar time dynamics to a simulation (**Fig. 2C**, top). This polarity in voltage at a particular sEEG electrode may be due to bipolar re-referencing and/or a nearby source of the electrophysiological activity. **Figure 3A,C,E** show the cortical distribution, magnitude, and statistical significance of *r*-based similarity values.

After permutation testing, an sEEG electrode was assigned significance if it had an FDR-corrected p-value < 0.001 for at least one of the three models. Out of 1258 total electrodes, 733 showed top-down similarity, 649 showed bottom-up similarity, and 634 showed SAM_y_ similarity. These include electrodes that were statistically significant for more than one model. All electrodes, regardless of significance, were included in further analysis.

Across all electrodes, similarity metrics from the three models and the control condition were significantly different in an omnibus analysis with a linear mixed-effects (LME) model fit to similarity, quantified as |median(Pearson *r*)|, with model type as a fixed effect and participant as a random intercept. Post-hoc analysis of estimated marginal means revealed that, compared to the audio envelope control condition, sEEG was more similar to the top-down model and the SAM_y_ model, while the bottom-up model’s similarity to sEEG fell below the control. Comparing between models, sEEG was most similar to the SAM_y_ simulation, followed by top-down, then the bottom-up model (**Fig. 3B,D,F; Table 3, Supp. Table 1**).

Because of individual differences in anatomy and electrode placement, fine cortical parcellation and dense anatomical labeling was not used in this analysis. Instead, electrodes were assigned to ten coarse anatomical regions, based on groupings of the Desikan cortical atlas^35^ (see **Table 2** for a summary of cortical parcels included in each region). However, all participants had coverage in the superior temporal cortex and prefrontal cortex. Regional differences in model similarity were analyzed with an LME model, similar to the whole brain approach, but with region interacting with model type.

The strongest model similarity effects are seen in the superior temporal cortex, insula, lateral temporal cortex, and prefrontal cortex (**Fig. 3A-G**, **Table 3)**. The cortical distribution of sEEG similarity to the presented models extended beyond the auditory evoked response (**Supp. Fig. 4**). Comparing models within regions, a familiar hierarchical gradient emerges across the cortex. Neural engagement in attentional modulation – via the top-down model – and integrative sensory anticipation – via SAM_y_ and the auditory envelope – are concurrent and spatially distributed, but not isolated. In early sensory regions like the superior temporal cortex, SAM_y_ and the sound envelope dominate – demonstrating the co-presence of the incoming signal and excitatory ramping activity anticipating each sensory input. Proceeding into the parietal cortex, a region associated with time estimation and auditory prediction^36,37^, this ramping effect surpasses the representation of the sound envelope, as SAM_y_ exceeds all other models. Top-down attentional processes contribute more in higher order processing regions in the lateral temporal cortex, where the top-down model and SAM_y_ are not significantly different. The top-down model surpasses SAM_y_ in the prefrontal cortex, reflecting neural engagement in attentional drive beyond sensory processing^17,38^.

This spatially-dualistic relationship between attentional bias and sensory input recapitulates accepted findings about simultaneous, complementary processes. It also highlights how well dynamical systems models of sensory anticipation capture neural activity in temporal and parietal regions. Most notable, however, is that our simulation approach can reveal this foundational structure while remaining within the bounds of plausible physiology and the cognitive demands of the temporal expectation task. Remarkably, these results are obtained without transforming the neural signal.

### Anticipatory activity in multiple regions predicts trial-wise subjective perception

Many electrodes exhibited some resemblance to the simulated LFPs, but merely resembling a model on its own does not indicate a meaningful relationship to the cognitive processes underlying auditory temporal expectation. To begin to address this, we sought to answer the following questions: if an sEEG signal resembled a model on a trial, would a participant make a more accurate timing judgement of the target tone? If so, which model provided the strongest advantage? To investigate this, we performed a mixed-effects logistic regression where the binary dependent variable of behavioral accuracy – whether the target tone was correctly judged as early or late – was classified by a fixed effect of the interaction of model type (top-down, bottom-up, SAMy, and audio envelope) with the simulated LFP’s similarity to the sEEG on that trial (|Pearson *r*|). Participant was included as a random intercept to account for multiple observations per participant and individual differences in behavioral accuracy (**Supp. Fig. 1**).

We first performed this mixed-effects regression on the whole brain, using all artifact-free sEEG electrodes. The whole-brain regression captured a small amount of variance in behavioral accuracy (Tjur’s R^2^ = 0.06), and an estimated marginal means analysis of the interaction fixed effects revealed a significant positive slope of SAM_y_-sEEG similarity and significant negative slopes of the top-down-and the audio envelope-sEEG similarities (**Fig. 4A**).

To examine the spatial distribution of neurobehavioral relationships to accuracy, a logistic regression of accuracy from the interaction of model type and similarity value was fit to all trials at each sEEG electrode. Participant identity was not included as a random intercept because each electrode only pertained to one individual. While the electrode-wise mapping of Tjur’s R^2^ indicated neurobehavioral relationships were relatively global (**Fig. 4B**), meaningful differences in each model’s relevance to accuracy emerged when model outputs were grouped by region (**Fig. 4C**). Specifically, SAM_y_-sEEG similarity was positively associated with accuracy in the superior temporal cortex, parietal, and prefrontal regions, and the top-down model had significant negative slopes in the lateral temporal, parietal, prefrontal, and orbitofrontal regions.

A similar mixed effects-regression captured participant’s trial-wise reports of confidence, with a marginally stronger effect than accuracy (Tjur’s R^2^ = 0.11) (**Supp. Fig. 5**). However, no model’s similarity to sEEG positively predicted confidence. Instead, the top-down, bottom-up, and audio envelope had significant negative slopes, suggesting that resembling any of these three models was related to lower confidence reports. These results were strongest in sEEG contacts in frontal regions. SAM_y_-sEEG similarity showed no significant relationship to confidence in any region.

The consistent negative relationship of the top-down model to accuracy – and confidence – is notable, such that on trials when the neural signal most resembled a simulation of top-down attentional bias, the participant was less likely to accurately or confidently judge the target tone as early or late. One plausible interpretation is that the top-down model does not account for the actual, jittered timing of the target tone, only its anticipated timing (**Fig. 2A**). Relying too heavily on expected timing could bias excitatory neural gain to the wrong moment, leading to the auditory afferent signal arriving during an insufficiently sensitive state, akin to neural oscillation phase biases observed in studies of sensory perception^39^. From a theoretical perspective, resembling the top-down model could indicate overreliance on a Bayesian Prior, which becomes indistinguishable from a prediction when sensory input is not sufficiently accounted for^24^. In contrast, the sensory anticipation model ramps excitation more adaptively to the onset of the arriving stimulus, with higher cortical sensitivity at the actual timing.

## Discussion

In cognitive neuroscience, neural signals that are measured during behavior reflect the combined influence of many concurrent computations. We present a novel experimental and analytical approach to differentiate overlapping neural processes and apply it to study auditory temporal expectation. We designed a subjective auditory timing task that manipulated the temporal certainty of an anticipated acoustic stimulus in an isochronous sequence of tones. In humans performing this task, greater perceptual accuracy was observed at longer jitters and with a greater number of prior tones before the target, suggesting that participants were able to better learn the temporal interval with the accumulation of more information. Importantly, our task design was such that every trial has a different IOI and a different jitter for the final tone to reduce the contribution of learning intervals across the experiment. While traditional approaches to analyzing the data from such a task would rely on averaging across trials or transforming the signal to analyze pre-defined frequencies, we took the novel approach of simulating trial-wise neural field potentials predicted from spiking models of top-down bias and sensory input. This approach allowed us to investigate the multiple, concurrent neural mechanisms of temporal expectation behavior by directly comparing simulated field potentials to invasively-recorded signals from participants performing the task. We found that sEEG signals resemble simulations in cortical regions associated with auditory processing including the temporal, parietal and prefrontal cortices. Importantly, top-down bias and anticipatory sensory activity in predicted trial-wise accuracy on the task.

These observations from invasive recording in humans align with previous studies in animal models, which show that early sensory cortices mediate perceptual effects of temporal expectation^15,16,40^. Beyond regions associated with acoustic perception, the cortical representation of predicted field potentials in regions like superior frontal, inferior parietal, and superior temporal cortex support previous findings of networks for voluntary attentional control^41,42^ and beat perception^43,44^. The relevance of anticipatory cortical activity to subjective perception exemplifies the utility of recent models of sensory anticipation^25^ and highlights an opportunity to test similar dynamical systems models of excitatory/inhibitory interactions in decision-making and memory in human field potentials^45^.

Our work has demonstrated how computational models, informed by known neurophysiology, can be implemented through biophysical simulation to directly interrogate their mechanistic potential in human cognition. The analyses presented here juxtapose three qualitatively different models, each representing a different component of the presumed neurocognitive process of auditory temporal expectation. Despite their differences, each model represents a plausible mechanistic component that adaptively contributes to auditory timing behavior, and all three likely occur simultaneously. Here, we provide a method to identify and describe each process in human participants, without transforming neural activity measurements based on assumptions about what is or is not relevant in a neural signal. Instead, these *a priori* assumptions are isolated to the selection of models and generation of the simulated LFPs. In some ways, the approach that we used resembles existing forward-encoding models of temporal response fields (TRFs), which describe how sensory information is encoded in neuronal activity as a function of time^46–48^, but unlike TRFs, our simulations do not require training data to estimate the predicted neural response.

The success of the sensory anticipation model in representing neural activity and predicting subjective behavior is notable but doesn’t preclude the relevance of other temporal prediction models. Future work could leverage our simulation approach to compare such dynamic models with other types in the temporal prediction space, including oscillatory models^13^, and timekeeper models^49^ to identify which of these models best predicts LFP results in which temporal prediction tasks. Notably, the bottom-up model never surpassed the audio amplitude envelope in its similarity to sEEG signals. Given that the spiking model employed in the bottom-up LFP simulation characterizes physiological responses in the cochlear nerve – a low-level response that is several synapses away from the primary auditory cortex – this bottom-up simulation may not best capture cortical dynamics. Future work can explore alternative bottom-up models of cortical responses to auditory input. Beyond model comparison, simulated signals could be applied to cortical surface electrodes or further forward-modeled to generate hypothetical non-invasive measurements from source-localized EEG and magnetoencephalography.

A limitation in the study is that the biophysics used in the simulations were highly simplified, with a single excitatory postsynaptic voltage kernel being convolved with the spike models. sEEG likely records an LFP comprising several different transmembrane currents^27^, and the resulting extracellular potentials also depend on the laminar location of the recording electrode^50^. Future applications of this approach can employ more complex and biophysically realistic models, including circuit dynamics^51^. It is notable, however, that in our sensitivity analysis of the effect of the synaptic kernel timescale, increasing the timescale substantially affected the correlation of the top-down and bottom-up models with sEEG (**Supp. Fig. 8**). The observation that different synaptic time constants yield different results implies that this biologically inspired modeling approach has the potential for meaningful application in translating theoretical spiking models into testable hypotheses in the LFP given more physiologically detailed models of synaptic distributions. It also highlights an opportunity to extend this correlational approach to controlled experimentation, wherein simulated LFPs could be validated against pharmacological manipulations of synaptic activity in non-human models. That said, the simulations that we used here represented an idealized process, as each spike field model was based on exogenous stimulus parameters.

Beyond the direct interpretations of the results obtained from this experiment, we believe that our modeling approach contributes a novel and flexible tool to our arsenal in the effort to combine advances in cellular and systems neuroscience with cognitive theories of behavior in humans. The approach that we employed here provides an interdisciplinary solution to some of the problems posed by the spatial limitations of non-invasive human recording^52^, the hazards of trial-averaging^53,54^, and the assumptions of periodicity and sinusoidality in the analysis of neural population activity and field potentials^32,53,55^. By leveraging recent advances in biophysical simulation^56^ and applied computational models of cognition^57^, we can expand the use of testable dynamic models of large scale brain activity^58,59^ to humans. Such an approach can help bridge scales between the mechanistic understanding of neural circuits and systems derived from model organisms to human cognition.

## Methods

### Neural data simulation

*Local Field Potential Simulations* Local field potentials were simulated to represent hypothesized neural activity in the auditory cortex from three different theoretical models: (1) a model of top-down bias, or dynamic attention^33^, (2) a model of (bottom-up) sensory inputs^7^, and (3) a model of dynamic sensory anticipation^25^. Each model was used to generate a function representing the probability of a postsynaptic current occurring in auditory regions on a trial-by-trial basis in the auditory attention task, taking into account the trial’s number of tones, IOI, and jitter.

Hypothetical LFPs were generated by summating simulated postsynaptic currents from a population of 100 spiking neurons. In this simulation, a uniform spiking population of 100 neurons were interpolated with a function representing hypothesized spiking probability for each signal type in each model where the probability of a spike occurring at any moment ranged between [0.05, 0.5]. Once interpolated with the probability function, spikes were convolved with a 50ms double exponential kernel representing the extracellular postsynaptic current of an excitatory glutamatergic (AMPA) receptor.

Postsynaptic AMPA currents were represented by a double exponential kernel described by:

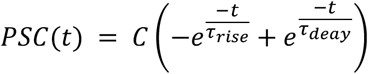

Where C is an amplitude normalization constant. For AMPA currents, τ_rise_ = 0.1ms and τ_decay_ = 2ms [CNRGlab @ UWaterloo].

*Bottom-up sensory input model* A bottom-up auditory sensory probability function was represented as a sequence of auditory nerve responses, as modeled by Huet et al., 2022^7^. Each response is characterized by an instantaneous increase in spiking (*a_r_*) at the onset of the stimulus, followed by an exponential decay (*τ_r_*), then a less extreme short term response and exponential decay (*a_st_*, *τ_st_*). A plateau (*a_base_*) in spiking probability continues until the end of the tone. This response repeats for each tone in the sequence. This function can be described as follows:

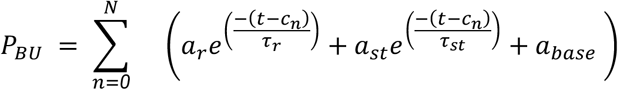

*Top-down bias model* A top-down attentional bias probability function was represented as a sequence of narrowing gaussians, beginning with the onset of the 2nd tone (the first prediction) and whose peaks are synchronized to the onset (*c*) of each tone. With each subsequent inter-onset interval (*b*) and tone (*n*) in the trial, the width (w) of the synchronized gaussian narrows, representing a learning process (*l*) which increases precision of the prediction of the arriving stimulus. Furthermore, the height (*a*) of the gaussian increases with each subsequent tone in the trial, representing a sensitization (*s*) for each expected tone. This function can be described as follows:

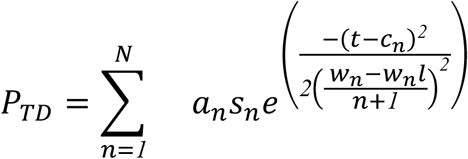

Parameter settings for both the bottom-up model and the top-down model are available in the supplement.

Spikes interpolated with top-down and bottom-up probability functions were convolved with double exponential kernels representing excitatory postsynaptic AMPA currents. The converging model describes theoretical modulations of neural excitability in expectation of an arriving stimulus and AMPA is the most common excitatory neurotransmitter.

*Sensory anticipation model* Local field potentials were simulated to represent the activity of a theoretical neural circuit for human sensorimotor timing described in Egger, Le, & Jazayeri, 2020^25^, from which our methods are borrowed. Specifically, this analysis employed the sensory anticipation module (SAM) of the full circuit model, which predictively ramps neural activity such that the output of the circuit reaches an expected level at the anticipated time of the next event.

The SAM spike module consists of four units that represent the average firing rate across a population of neurons. Here, we use the same naming convention as in the original paper, such that the four units are *I*, *u*, *v*, and *y*. *u* and *v* are mutually inhibitory, and both receive input, *I*, that is tonic and remains constant over time until it is adjusted. The output of the model, *y*, receives excitatory input from *u* and inhibitory input from *v*. The mutual inhibition between *u* and *v* results in the ramp-like activity that is essential for the sensory anticipation dynamics. The implementation of this anticipation relies on an error signal, *y*-*y0*. The model adjusts its input (*I*) such that *y* reaches *y0* at the moment when the tone is expected, at a rate proportional to the error signal. This module evolves according to the following equations:

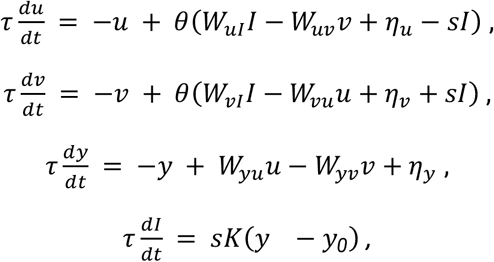

Here, τ is the time constant for each unit, θ(x) is a sigmoid activation function (*θ*(*x*) = 1/[1+exp(−*x*)]), and *I* is a tonic input. The weights for all the unit interactions, W_uI_, W_uv_, W_vu_, and W_vI_ were set to 6 for all simulations, while both W_yu_ and W_yv_ were set to 1, per the original implementation. Zero mean Gaussian noise (η) was added to units *u*, *v*, and *y*, with variance σ. *K* is a free parameter. Input (*I*) is set to 50, and given a binary weight *s* that is equal to 0 when the stimulus is absent, and 1 when the stimulus is present.

Sequences of stimuli were implemented for each trial by setting s=1 at the onset of each tone in the trial for 10ms. All other model parameters for all simulations are defined in **Supp. Table 2.** In this analysis, the response of the *y* unit of the SAM was used as the spiking probability signal to simulate an LFP, and this model is referred to as SAM_y_ in the text. In supplementary analyses, simulations were also generated with the *u* and *v* units (See **Supp. Fig. 6**).

### Participants

#### Healthy controls

Eleven healthy control participants were recruited from the undergraduate student population at UC San Diego. All participants gave fully informed written consent, as monitored by the Institutional Review Board.

#### Intracranial participants

Data from ten patients (4 female, 29.2 ± 6.5 years old) with pharmaco-resistant epilepsy undergoing intracranial sEEG recording for seizure onset localization preceding surgical treatment were included in this study (**Table 1**). Participants included in the study had electrodes implanted in what was eventually found to be non-lesional, non-epileptogenic cortex (such areas were suspected to be part of the focus before implantation or were necessary to pass through to reach suspected epileptogenic areas). All participants gave fully informed written consent for their data to be used for research as monitored by the local Institutional Review Boards at UC San Diego Health.

### Behavioral task

Participants performed a novel behavioral paradigm designed to manipulate the temporal certainty about the timing of an anticipated stimulus by controlling the number of prior stimuli in a rhythmic stream (**Fig. 1A**). In this task, participants listened to a sequence of three to five 200 ms piano tones (C_4_–B_4_, 261-494 Hz) presented at a frequency of 1-3 Hz, or with an inter-onset interval (IOI) 333-1000 ms. All tones were presented isochronously (equally spaced in time), except for the last tone in the sequence, which was jittered based on a uniform distribution between ±0.4 times the inter-stimulus interval for any given trial. Participants reported whether the last tone in the sequence, hereafter referred to as the target tone, occurred earlier or later than they expected, responding with a key press of 1, 2, 3, or 4. A response of 1 indicated a confident report of an early target tone and 4 indicates a confident report of a late target tone. A response of 2 and 3 indicated less confident responses and participants were instructed to guess if they were uncertain. On any given trial, participants were not told whether the target tone would arrive at the third, fourth, or fifth position in the sequence. The participants were discouraged from moving (e.g., tapping) to the beat of the tone sequence to minimize motor artifacts and/or direct recruitment of motor regions. Participants reported age, sex, and any musical training to track any effects of musical expertise in rhythmic discrimination precision.

### Data acquisition

Experimental instructions and stimuli were presented to participants in their hospital rooms on a Windows 10 desktop PC (Dell XPS 8910) using PsychoPy v.3.1.0. Auditory stimuli were presented with a low latency audio soundcard (Native Instruments) fed into desktop speakers placed two feet from the patient at volume the patient indicated was clearly audible, but not uncomfortable. After an instructional block with nine practice trials, the task was conducted in four blocks, each of which contained 60 trials (249 trials total). All participants completed the full experiment.

sEEG signals were amplified using a multi-channel amplifier system (Natus Quantum) and recorded using Natus Neuro-Works software. Auditory stimuli were recorded simultaneously with the sEEG data by feeding the output of the audio soundcard as an additional input channel to the Natus Quantum amplifier. sEEG signals were sampled at 1024 Hz, except for one participant whose sEEG was sampled at 512 Hz. Sampling rate had no apparent impact on results.

After recording, neural data were de-identified and exported from the clinical NeuroWorks system in.edf (European Data Format) format. Prior to analysis, files were assembled in the iEEG-BIDS format using mne-bids in Python^60^.

### Data analysis

*Behavioral data analysis* Behavioral performance was measured using the slope of a sigmoid function fit to the performance of a participant making a perceptual judgment about the target tone:

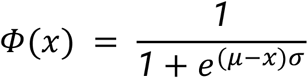

where **Φ** represents the probability of a ‘LATE’ response at a given jitter (*x*), measured relative to the IOI. The sigmoid function was fit to the average of binarized response (1 or 2 = EARLY, 3 or 4 = LATE) binned by jitter in 20 equal bins within the experimental range of ± 0.4 times the IOI. When fit to the behavioral responses of a participant performing the task, *μ* is the *equivalence point*, or the value of the jitter where a participant is equally likely to judge a target tone as late or early. Variations in equivalence point indicate a participant is biased to respond that an irregular tone is more often early than late or vice versa. *σ* is the *slope* of the sigmoid representing the participant’s perceptual accuracy. More accurate judgements of target tone irregularities will produce steeper sigmoid slopes.

*Electrode localization* sEEG electrodes were localized as per Marsh et al. 2024^61^. Briefly, each patient’s pre-op T1-weighted MRI and post-implant CT volumes were co-registered in 3DSlicer^62^ and CT artifacts indicating electrode positions were manually annotated. Cortical surfaces were reconstructed from the MR volume using the standard FreeSurfer recon-all pipeline^63^. Automated parcellation^64^ was used to assign atlas-based anatomic labels to regions of the cortical surface in the Desikan atlas^65^. A coarser parcellation into ten cortical regions of interest was obtained by grouping together regions from the Desikan atlas^35^ (see **Table 2** for groupings). All electrode locations on native pial surfaces are presented in **Supp. Fig. 9**. For visualization purposes, cortical surfaces and electrode locations were morphed from individual surfaces to the Freesurfer fsaverage space.

#### sEEG preprocessing

sEEG recordings were converted to iEEG-BIDS format^60^ prior to preprocessing. Only data from sEEG contacts were included in the analysis. Cortical surface electrodes from electrocorticography (ECoG) grids and strips were excluded for generalizability, as not all participants had surface coverage. Data from sEEG contacts were bipolar re-referenced and notch filtered at 60, 120, 180, and 240 Hz to remove line noise. Channels were excluded from analysis if they contained interictal activity, based on clinical indication. On average, 27±14 channels were excluded per patient. sEEG data was epoched into 5.8s epochs, beginning 0.8s prior to the presentation of the first stimulus in each trial. Epochs were visually inspected and epochs containing interictal activity that spread beyond the clinically-identified channels were excluded from analysis. On average, 19±22 epochs were excluded per patient. **Table 1** contains information for channel and epoch exclusions for each patient. Prior to any further analysis, data were resampled to 1000 Hz.

#### Simulation comparison to sEEG

To determine how strongly LFP simulations from the converging inputs model and the sensory anticipation model resembled the neural activity of participants performing the temporal expectation task, simulated LFPs were compared with sEEG signals. For each trial, the hypothetical LFPs of each model were simulated for the entire length of the 5.8s epoch. For consistency across experimental conditions, a segment beginning 700ms before the onset of the first tone to 400ms after the last tone was extracted for all simulated LFPs and sEEG data. The LFPs and the sEEG data for each trial were resampled to a fixed length of 2500 samples to mitigate the potential effects of signal length on correlation strength. For each trial, all three simulated LFPs (bottom-up, top-down, and sensory anticipation) were correlated with the sEEG signal using a Pearson correlation. The median correlation coefficient, Pearson’s *r*, across trials was computed for each sEEG channel, representing how well that channel resembled the simulation. Absolute value was computed after finding the median because the *r*s from a given sEEG electrode were normally distributed across included trials. Taking the absolute value of all *r*s prior to computing a median would generate a non-normal distribution, for which a median is not an appropriate measure of central tendency.

#### Control analysis

As a control analysis, we correlated the amplitude envelope of the presented auditory stimulus to the LFP on each trial. The auditory signal was collected as a digital copy of the auditory stimulus input to the amplifier from the external sound card via an auxiliary splitter cable. The audio signal was sampled at the acquisition rate of the amplifier (1024 Hz) and resampled at 1000 Hz. On each trial, the amplitude envelope was computed as the absolute value of the Hilbert transform of the bandpass filtered signal between 260-495 Hz. This frequency range was selected to extract the fundamental frequency of the piano tones used in the task (C4-B4, 261-494 Hz). For smoothing, a 21-sample median filter was applied to the amplitude envelope. Example auditory amplitude envelopes are depicted in **Supp. Fig. 7**.

#### Software

The behavioral task was presented using PsychoPy v3.9. sEEG preprocessing was performed using mne-bids v0.15.0 and MNE v3.7.0. LFP simulations were performed using neurodsp v2.2.1 and custom functions in Python. Other analyses were performed with custom code in python v3.10.8 using numpy v1.21.6, scipy v1.10.0, pandas v1.5.2. Statistical analyses were performed using pingouin v0.5.5, statsmodels v0.14.4 and scikit-learn v1.6.0. Mixed-effects models were implemented in R with lme4 and emmeans.

## Statistical analysis

*Within-participants behavior* Differences in perceptual accuracy of precise timing judgments between experimental conditions in the temporal expectation task were assessed by performing a one-way, repeated-measures ANOVA on the equivalence point (*μ*) and slope (*σ*) values extracted from a sigmoidal function fit to each participant’s performance. The within-subject factor of the experimental condition (number of tones) was analyzed for its effect on the dependent variable (slope or equivalence point) across participants. Significance was computed as a p-value adjusted for violations of sphericity using a Greenhouse-Geisser correction when appropriate. Post-hoc repeated-measures, two-tailed t-tests were used to assess differences in equivalence point and slope between experimental conditions. Differences in metrics between conditions were considered significant if the p-value < 0.05 after adjusting for multiple comparisons with a 6-test Holm-Bonferroni correction.

*Between-participants behavior* Potential differences in performance between healthy controls and sEEG participants included in the study were assessed by performing a one-way mixed ANOVA. The within-subject factor of experimental condition (number of tones) and the between-subject factor of experimental group (sEEG or healthy control) were analyzed for their effects on the dependent variable (slope or equivalence point) across participants. Post-hoc independent-measures, two-tailed Welch’s t-tests were used to assess differences in equivalence point and slope between experimental groups (sEEG or healthy control) in each experimental condition (number of tones). Differences in metrics between groups were considered significant if the p-value < 0.05 after adjusting for multiple comparisons with a 6-test Holm-Bonferroni correction.

*Permutation Testing* The significance of electrode-wise correlations between simulations and sEEG data were tested using a 1000-fold permutation test. In each permutation, trial parameters (trial IOI, jitter, and number of tones) were randomly shuffled across all included trials and then were used to generate simulated LFPs for each model. Simulated LFPs were compared with the sEEG signal on that trial using Spearman correlation. The segment of sEEG data remained the same as the original analysis (700ms before the onset of the first tone to 400ms after the last tone), and the simulated LFP signal was cropped to match the length of the sEEG data to maintain identical timescales and signal lengths. In each permutation, the median Pearson *r* for each channel was computed across all included trials, producing a distribution of 1000 values. P-values were computed akin to a one-sided t-test. If a sample Pearson *r* was negative, its p-value indicated the proportion of the distribution that was more negative than the sample, and vice versa for positive values of Pearson *r*. A Pearson *r* for an electrode was considered significant if its p-value < 0.001 after FDR correction with a Benjamini-Hochberg procedure (scipy.stats.false_discovery_correction). Only significant electrodes were included in further analysis.

*Comparison of sEEG-simulation similarity between models.* Linear mixed-effects models were used to compare sEEG similarity between models where the similarity of a model at a given electrode was classified by a combination of fixed effect of model type with participant as a random intercept:

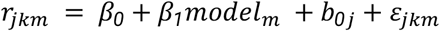

where *r_jkm_* is the summarized similarity metric |median(Pearson *r*)| across all trials for electrode *k* from participant *j* for model *m*, and

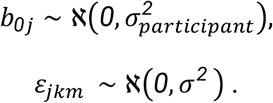

Where *β_0_* is the fixed intercept, *β_1_* is the fixed main effect, *b_0j_* is the random intercept, and *ɛ_jkm_* is the residual error. Statistical significance and pairwise contrasts of the fixed effects were assessed using Wald-z tests of estimated marginal means and p-values were FDR-adjusted for multiple comparisons.

To compare model similarity within regions, a similar mixed effects model was used, with the added interaction effect of cortical region, based on the electrode’s native location in a coarse grouping of the Desikan atlas.

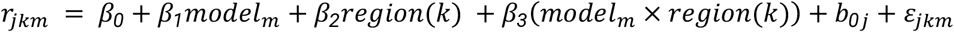

where region is an attribute of an electrode *k*. *β_0_* is the fixed intercept, *β_1_* and *β_2_* are the fixed main effects, *β_3_* is the fixed interaction effect, *b_0j_* is the random intercept, and *ɛ_jkm_* is the residual error. For interactions between model type and region, conditional model effects were estimated within each anatomical region using the emtrends function from emmeans in R. Pairwise comparisons between models were then performed within each region using Wald z-tests, and p-values were FDR-adjusted for multiple comparisons.

## Statistical modeling

*Mixed modeling of behavior & sEEG* We sought to investigate the hypothesis that an sEEG signal that resembles one or more of the proposed spike models is related to behavioral accuracy on the auditory timing judgment task. To do so, we used a generalized LME model with a binomial link function, where a binary variable of judgment accuracy (correct or incorrect late/early judgment) on a trial was classified by a combination of fixed and random effects. The interaction of similarity (|Pearson *r*|) with model type (top-down, bottom-up, SAM_y_, and audio envelope) was included as a fixed effect, and participant identity was treated as a random effect, as individuals varied in their behavioral performance on the task (**Supp. Fig. 1**). This model was fit to all trials and all electrodes, pooled across participants. Stimulus features like IOI, number of tones and jitter used to derive the simulated models, so they were not included in the multiple regression. For a single trial, this relationship is described by the formula:

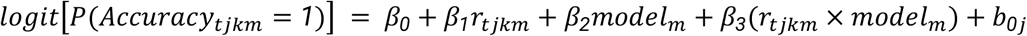

where *r_tjkm_*is the similarity metric |Pearson *r*| on trial *t* for electrode *k* from participant *j* for model *m*, and

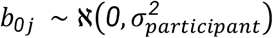

Where *β_0_* is the fixed intercept, *β_1_* and *β_2_* are the fixed main effects, *β_3_* is the fixed interaction effect, and *b_0j_* is the random intercept. Conditional slopes associated with each model type were estimated from the fitted interaction model using the emtrends() function in the emmeans package in R. Reported p-values are extracted from Wald-z tests of the whole-brain estimates. Model explanatory power was computed with Tjur’s R^2^, a discrimination-based metric for logistic regression defined as the difference between the mean predicted probability for positive versus negative outcome classes.

For regional analyses, a similar multiple logistic regression was performed at each sEEG bipolar contact (electrode). Participant was not included as a pooling variable, as each electrode belonged to only one participant.

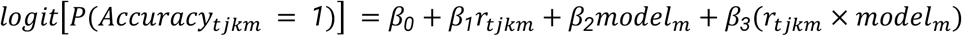

Electrode-level effect sizes were computed with Tjur’s R^2^. Conditional slopes associated with each model type estimated with emtrends() and were grouped by anatomical region prior to one-sample t-tests. Resulting p-values were FDR-corrected for multiple comparisons.

## Supplement

### Supplementary Methods

*LFP Simulation Sensitivity Analysis* To investigate the relevance of the postsynaptic voltage kernel used in the LFP simulation, we performed a parameter sweep of voltage decay timescale (τ_decay_) within a physiologically-plausible range of kernel lengths (2 - 200 ms). We also ran the analysis using only the spike probability signal (no kernel). Here the LFP simulation for all three spiking models was performed as described in **Methods**, with 500ms kernel lengths and τ_decay_ systematically varied, and outcomes were evaluated based on Spearman *ρ* coefficients and a single-variable logistic regression of behavioral accuracy (**Supp. Fig. 8**).

## Code & Data availability

All code used for all analyses and plots are publicly available on GitHub at https://github.com/voytekresearch/prophecy. Intracranial data will be made available upon reasonable request, with the proviso that sharing of raw data in particular is subject to approval by the Institutional Review Board.

## Acknowledgements

We would like to thank Fabian Schmidt for his valuable insights on our analysis.

Support: NIH National Institute of General Medical Sciences grant R01GM134363-01 (to B.V.) and NIH National Institute of Mental Health R61MH135109 (to B.V.).

## Author contributions

Based on CRediT roles

Conceptualization; S.E.S., K.B.D., B.V.

Data curation: S.E.S.

Formal analysis; S.E.S., A.R., K.B.D.

Funding acquisition; B.V., J.J.S.

Investigation: S.E.S.

Methodology; S.E.S., A.J.S, K.B.D., B.V.

Project administration; J.J.S., S.B.H.

Resources; S.B.H., J.J.S., B.Q.R., B.V.

Software; S.E.S., K.B.D., D.C., B.V.

Supervision; B.V., K.B.D., J.J.S., S.B.H

Visualization; S.E.S., J.C.G., B.Q.R.

Roles/Writing - original draft; S.E.S., B.V., K.B.D.

Writing - review & editing. All authors

## Ethics

Human subjects: All participants gave written informed consent to participate in the study. All experimental procedures were approved by the institutional review board of the University of California, San Diego, Human Research Protections Program (UCSD IRB Protocols #150834 and #161788).

## Supplementary Figures

**Supp. Fig. 1.**
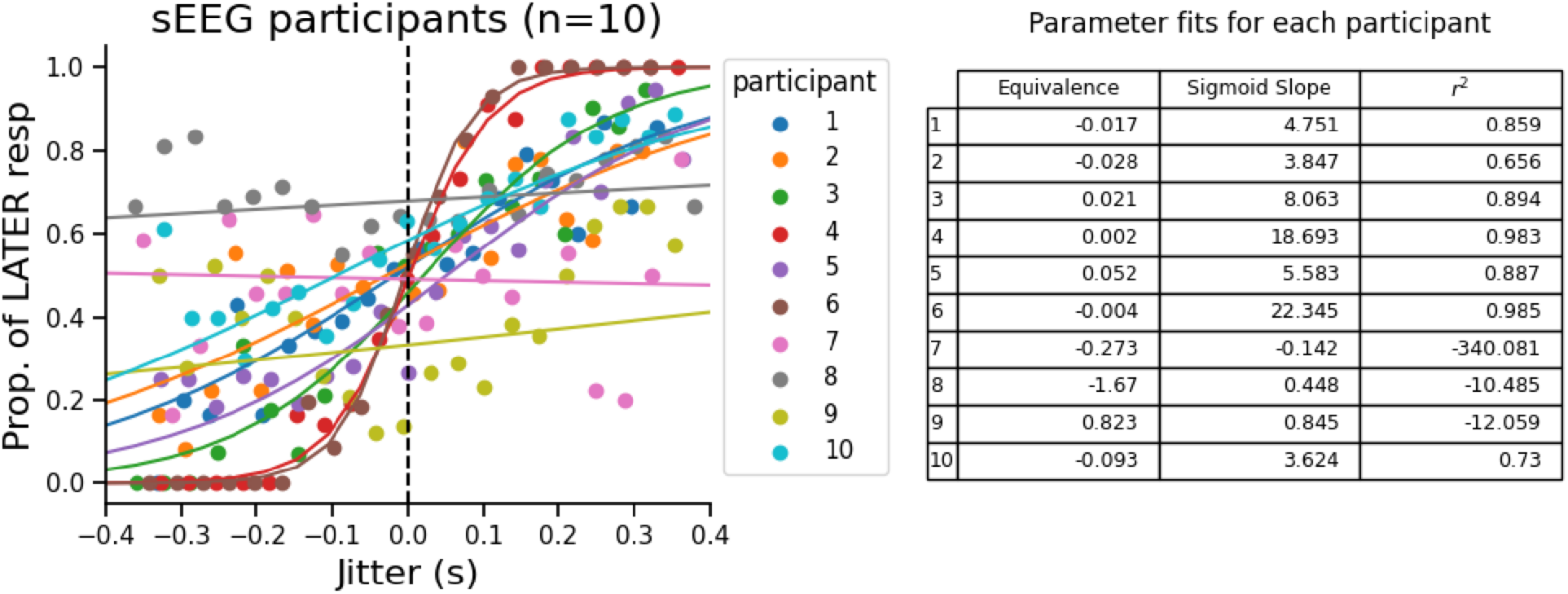
Individual differences in sEEG participant accuracy and engagement in task structure. Left: sEEG participant performance across the full task. Jitters (x-axis) and the proportion of LATE responses (y-axis) are averaged in each of 20 bins. **Right**: equivalence, slope, and weighted coefficient of determination score (r^2^) of the sigmoid fit to participant’s responses, left. Negative scores indicate the sigmoid model’s mean squared error is larger than the variance of the data.

**Supp. Fig. 2.**
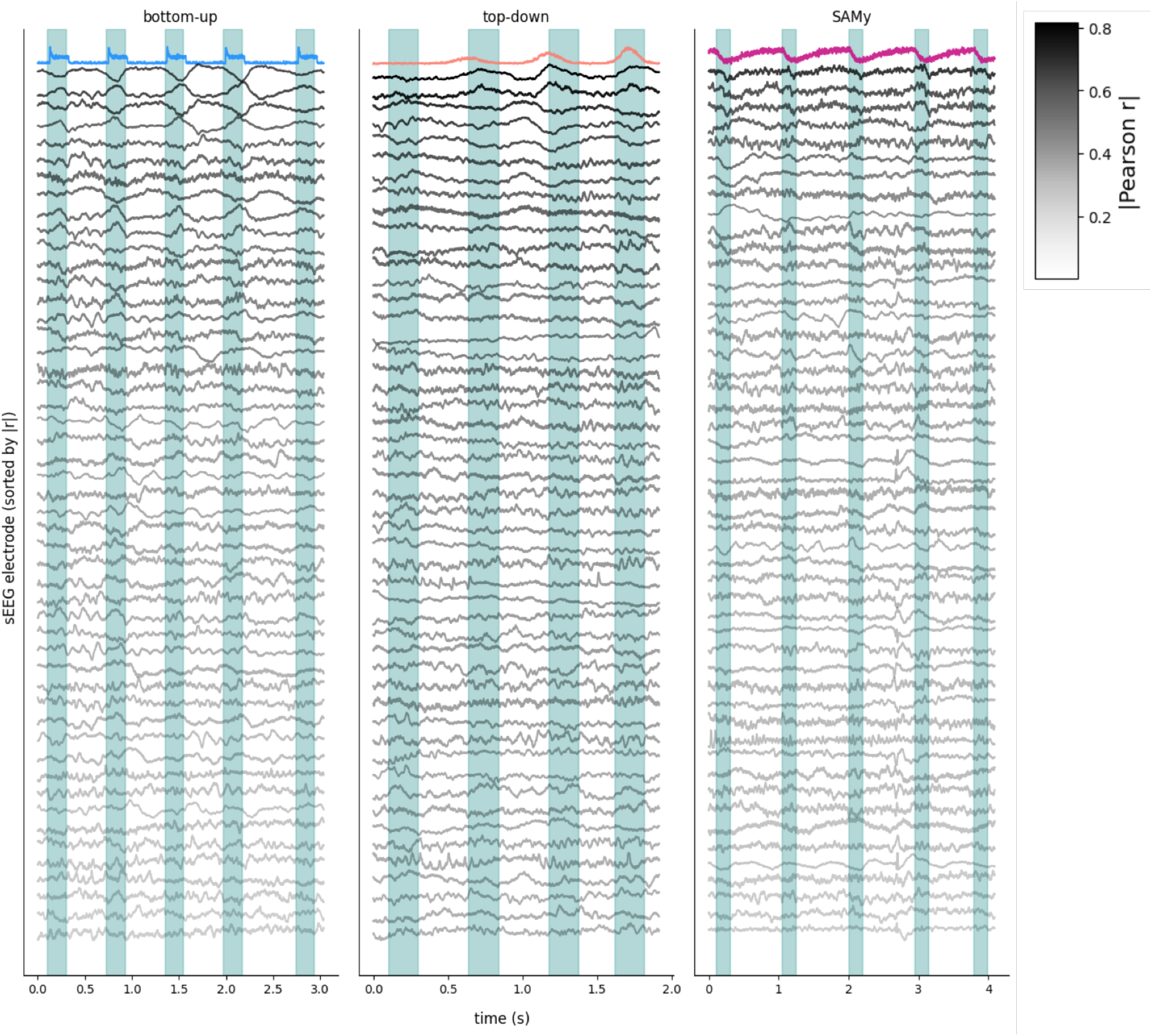
Inter-electrode variability of model similarity within trials. 50 sEEG electrodes from one participant in the three trials depicted in Figure 2C. Simulations are in color (blue: bottom-up, salmon: top-down, magenta: SAMy), sEEG data is in grayscale, where darkness represents higher similarity scores (|Pearson *r*|). sEEG electrodes are sorted by descending similarity, and are not the same electrodes in each column.

**Supp. Fig. 3.**
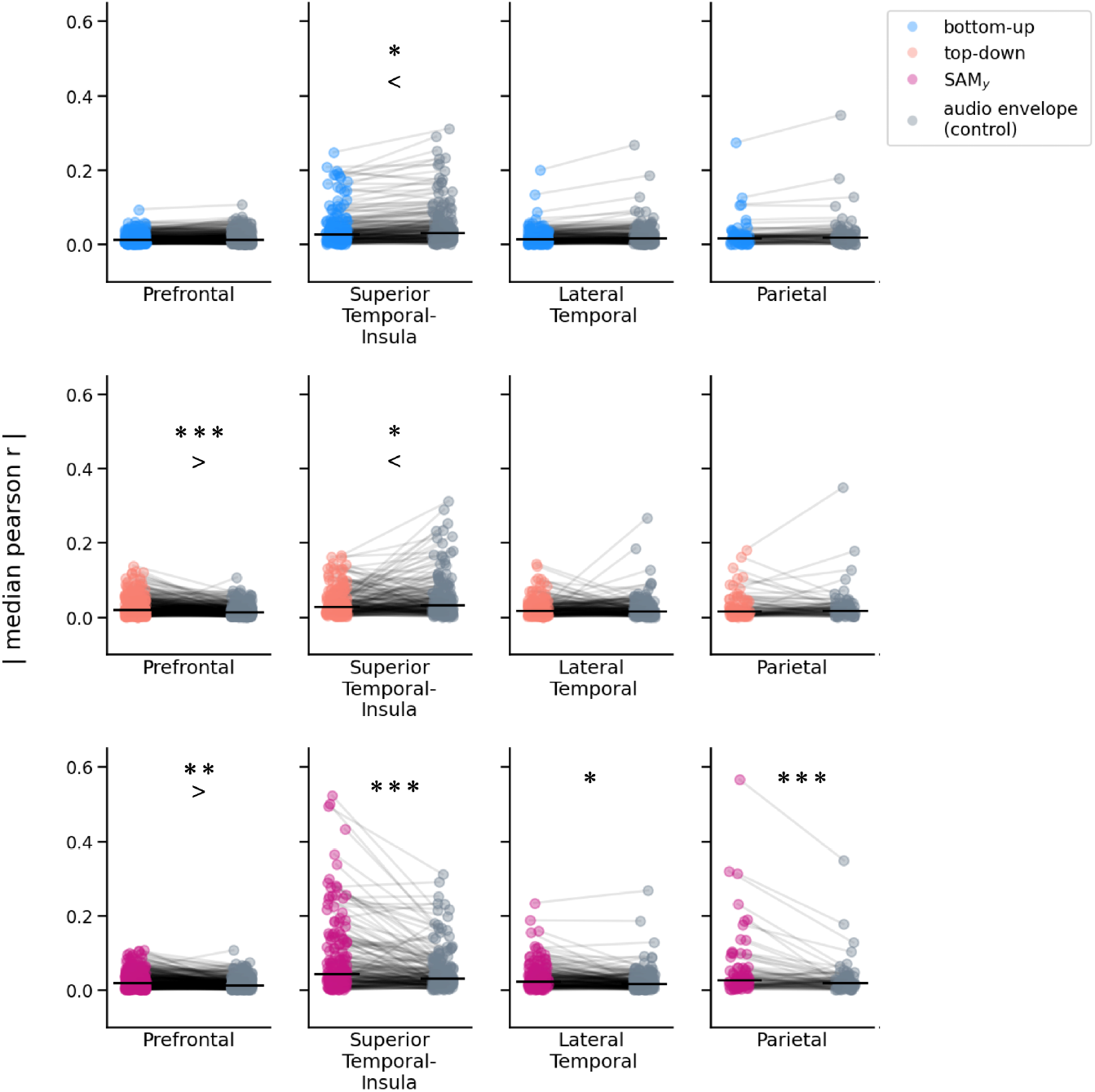
**Similarity values of experimental models and the control condition in electrode pairs**. An extension of the data displayed in Fig. 3D. Wald tests of estimated marginal means from a linear-mixed effects model identified significant differences between models and controls in four cortical regions: Prefrontal, Superior-Temporal cortex and Insula, Lateral Temporal cortex, and Parietal cortex. The audio envelope (control) condition-sEEG similarity surpassed the bottom-up-sEEG similarity and top-down-sEEG similarity in the superior temporal cortex. The top-down-sEEG similarity surpassed the control-sEEG similarity in the prefrontal cortex. The SAM_y_-sEEG similarity significantly exceeded the control-sEEG similarity in all four regions.

**Supp. Fig. 4.**
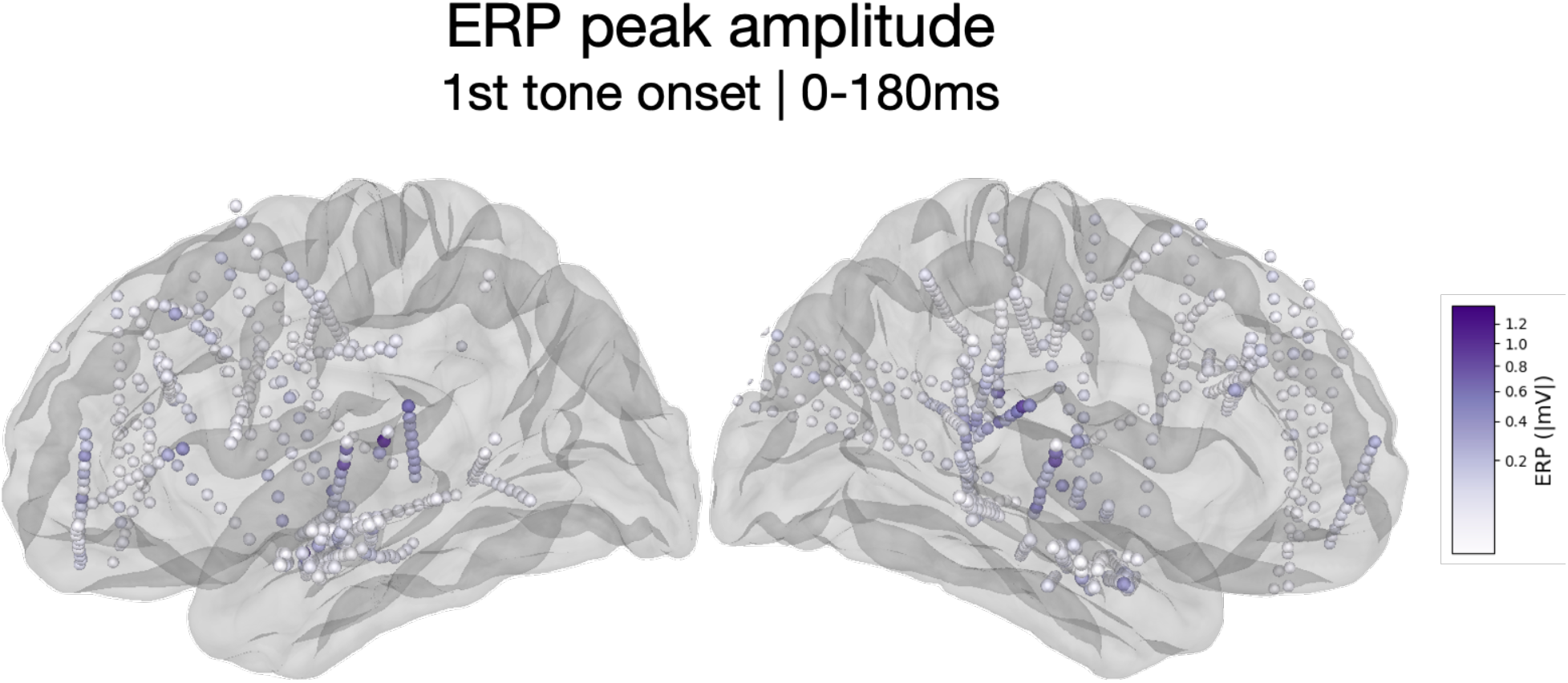
**Cortical distribution of the auditory evoked neural response**. The color of each sEEG electrode represents the maximum amplitude of the evoked response between 0-180ms after the first tone in the trial. ERPs were computed by downsampling the sEEG data to 250Hz and averaging over all included trials.

**Supp. Fig. 5.**
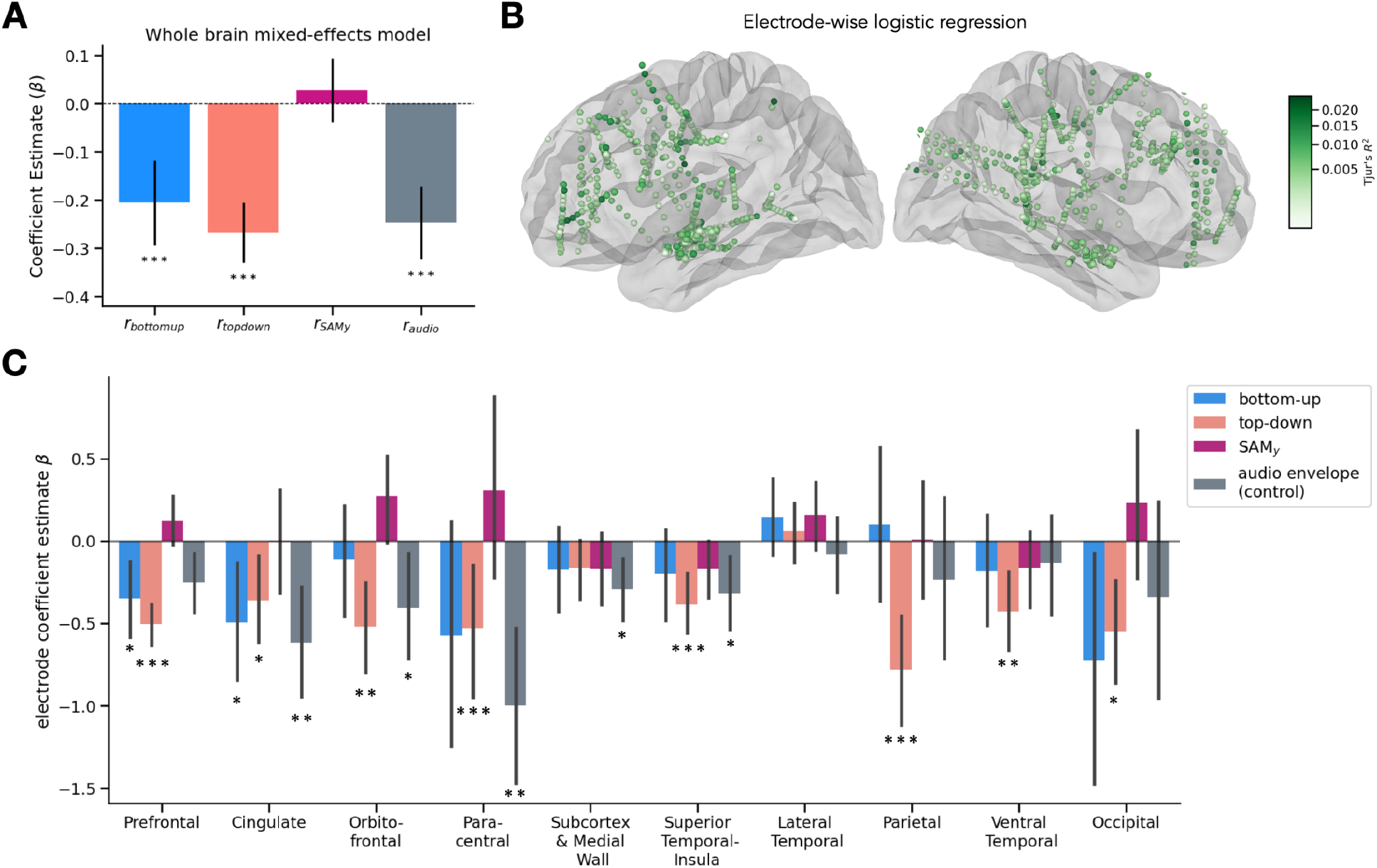
Mixed-effects modeling of trial-wise confidence from neural model similarity. **(A)** Estimated slopes (β) relating neural fit (r) to confidence report for each model type, derived from a generalized LME model with participant as a random intercept (Tjur’s R^2^ = 0.11). Bars indicate fixed-effect slopes from estimated marginal means; error bars represent 95% CI. The slope for the bottom-up model (β_bottom-up_ =-0.21, p < 10^-5^), top-down model (β_top-down_ =-0.27, p < 10^-5^), and audio envelope (β_audio_=-0.25, p = p < 10^-5^) were significantly negative. **(B)** Cortical map of electrode-wise logistic regression of accuracy. Electrode color represents Tjur’s R^2^. **(C)** Coefficients of electrode-wise regressions shown in panel B, distributed by cortical region. Bars indicate slopes from estimated marginal means; error bars represent 95% CI. Stars indicate significantly positive/negative slopes for electrodes in that region (one-sample t-test: ∗ p_FDR_ < 0.05, ∗∗ p_FDR_ < 0.01, ∗∗ p_FDR_ < 0.001).

**Supp. Fig. 6.**
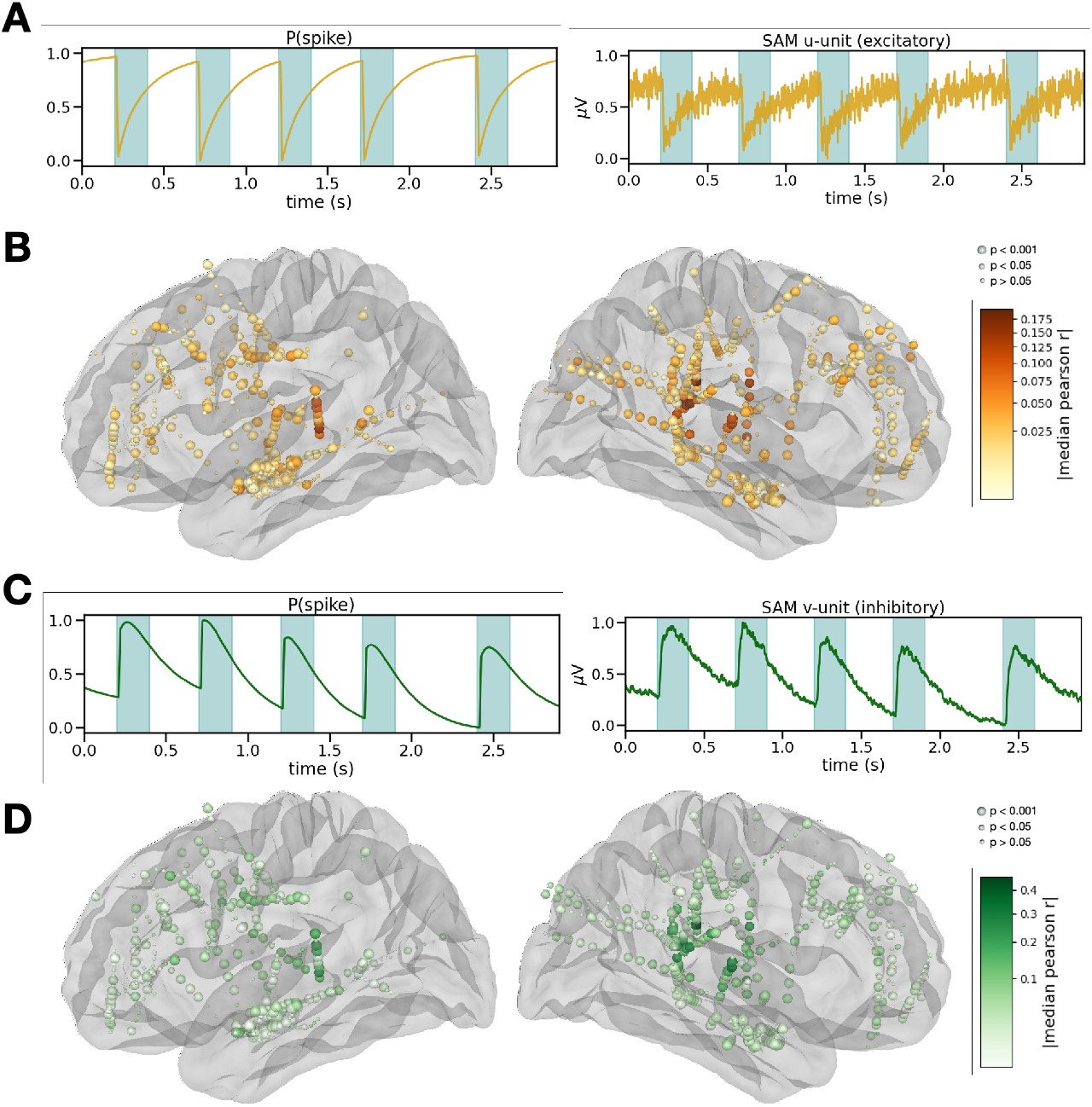
Sensory anticipation module (SAM) u-and v-units and sEEG similarity. **(A)** Left: Spiking probability signal of rate unit *u*, a unit with excitatory connections to the y-unit. Spiking probability plotted for an example trial (like Fig. 2) Right: Simulated LFP for the trial using an AMPA voltage kernel for spike convolution (τ_rise_ = 0.1 ms, τ_decay_ = 2 ms). **(B)** SAM (*u*-unit) model similarity to sEEG. Color intensity represents the |median(r)| at each sEEG electrode. Sensor size represents FDR-corrected p-value. **(C)** Left: Spiking probability signal of rate unit *v*, a unit with inhibitory connections to the *y*-unit. Spiking probability plotted for an example trial (like Fig. 2) Right: Simulated LFP for the trial using an GABA_A_ voltage kernel for spike convolution (τ_rise_ = 0.5 ms, τ_decay_ = 10 ms). **(D)** SAM (*v*-unit) model similarity to sEEG. Color intensity represents the |median(r)| at each sEEG electrode. Sensor size represents FDR-corrected p-value.

**Supp. Fig. 7.**
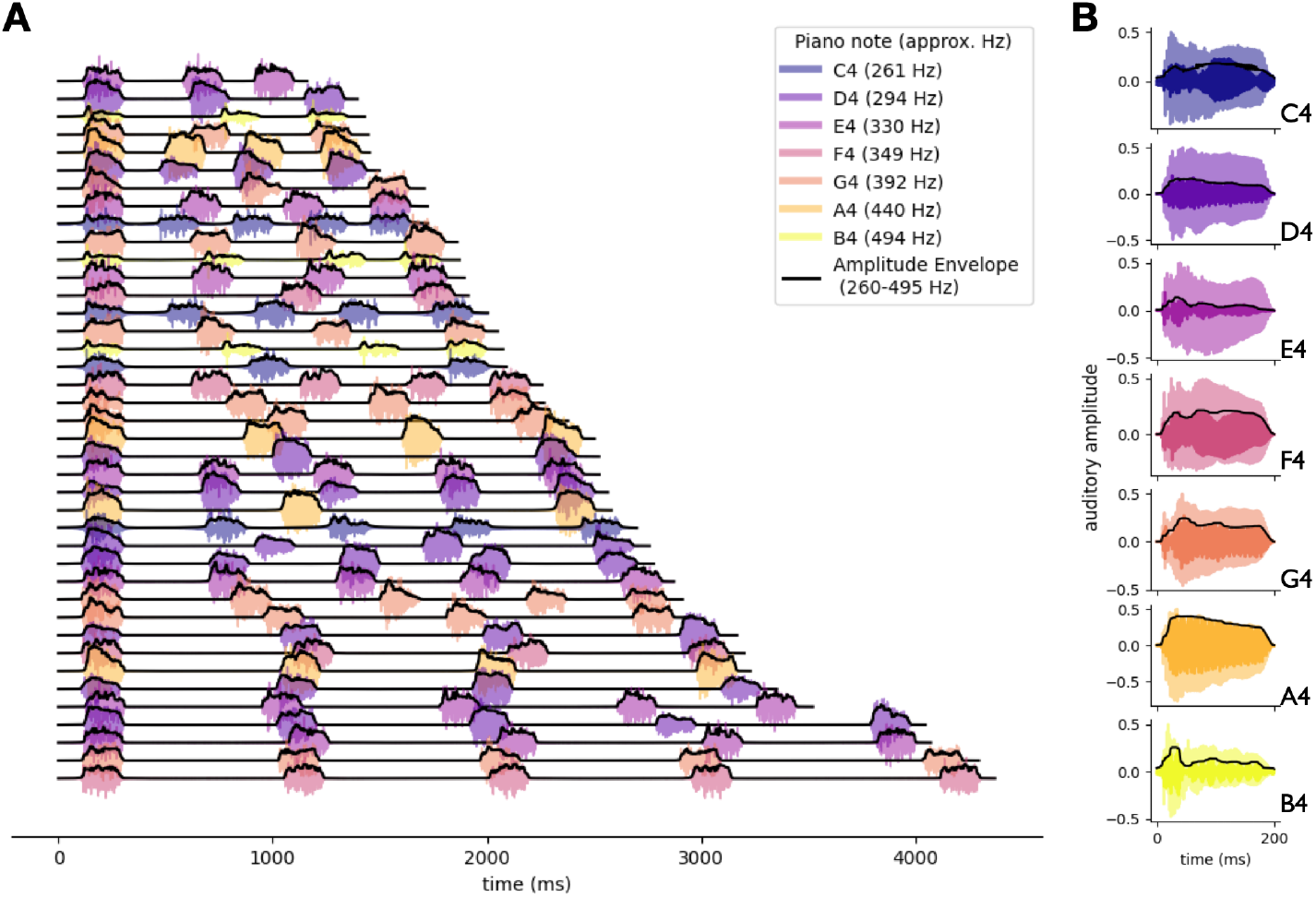
Audio amplitude envelope (control) example trials. **(A)** 40 randomly-selected trials depicting the audio amplitude envelope. Properties of the selected trials reflect the range used in the experiment. Each trial included three to five piano tones, IOIs between 333ms-1000ms, and jitters ±40% of the IOI. Each colored line is the amplitude of the audio signal, sampled at 1000 Hz and filtered between 260-495 Hz. Lines are color-coded based on the piano note played during the trial. Solid black lines denote amplitude envelope, computed as the absolute value of the Hilbert transform of the audio signal, smoothed with a 21-sample median filter. (**B)** Amplitude envelopes of single notes at original audio file sampling rate, 44100 Hz (translucent color), overlaid with the audio signal resampled at 1000 Hz and filtered in 260-495 Hz (solid color), and the smoothed amplitude envelope (black).

**Supp. Fig. 8.**
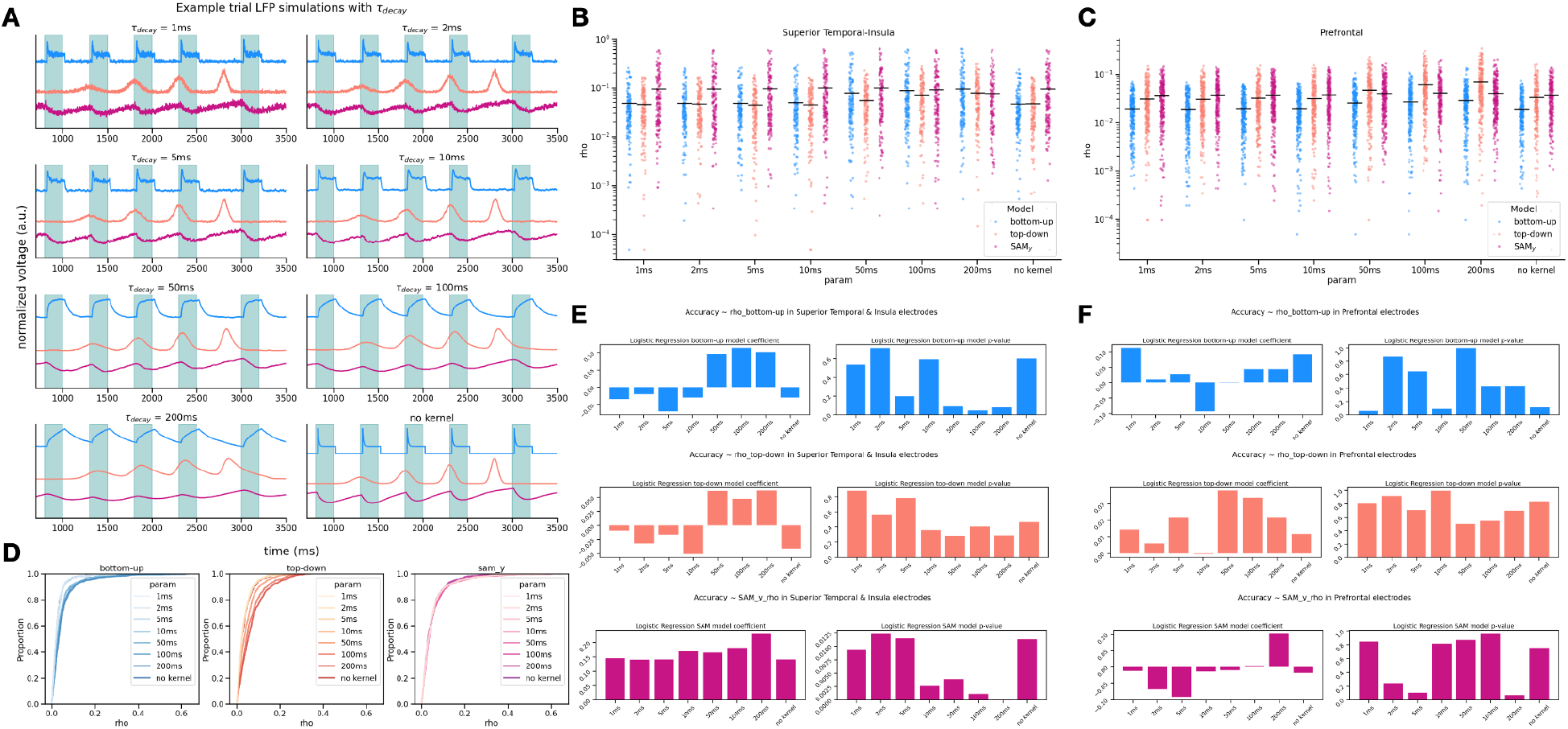
LFP simulation sensitivity analysis. **(A)** Example LFP simulation of a trial with five tones, an IOI of 500ms (2 Hz), and a jitter of 200ms. Each subpanel depicts the simulation of the bottom-up (blue), top-down (salmon), and SAM_y_ (magenta) spiking models with a systematically varied τ_decay_ in the convolved kernel representing the voltage trace of an EPSP. τ_decay_ = 2ms was used in Main. **(B)** Absolute value of the median Spearman ⍴ for sEEG bipolar midpoints in the superior temporal cortex and insula, electrodes are the same as those in Fig. 3D in Main. The x-axis represents the τ_decay_ of the simulated LFP. **(C)** Like panel B, but in the prefrontal cortex. **(D)** Empirical cumulative density functions for ⍴ across all trials and electrodes in sEEG participants. Bottom-up and top-down model rhos increase with greater τ_decay_, while the SAM_y_ model similarity is less dependent on τ_decay_. **(E)** Single-variable logistic regression coefficients and p-values from select sEEG electrodes in the superior temporal cortex and insula (see Methods). Here, only ⍴_SAMy_ is predictive of trial-wise accuracy, with a trend favoring greater τ_decay_. **(F)** Like panel E, but for sEEG electrodes in the prefrontal cortex. No model ⍴ significantly predicted trial-wise accuracy.

**Supp. Fig. 9.**
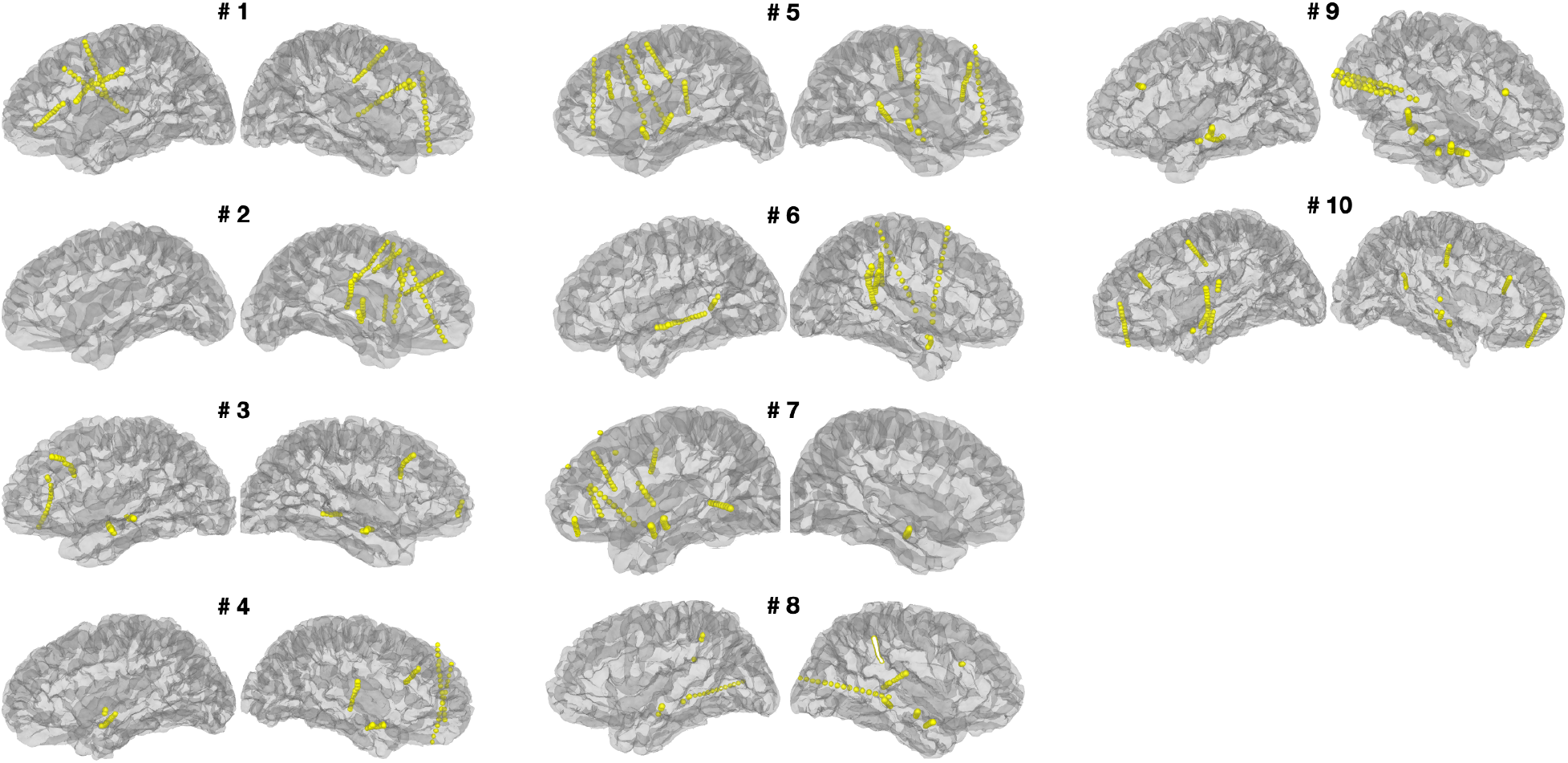
sEEG contact pair bipolar midpoint locations in native pial space. Electrode locations are presented on native pial surface for all seven intracranial participants. Participant IDs are the same as **Table 1**. See **Methods** for electrode localization procedures.

## Supplementary Tables

**Supp. Table 1.** Region-wise contrasts of estimated marginal means and FDR-corrected p-values.

| region | model 1 | model 2 | $\Delta r$ (EMM) | CI (5%) | CI (95%) | z | p(FDR) |
| --- | --- | --- | --- | --- | --- | --- | --- |
| Whole Brain | top-down | bottom-up | 0.007 | 0.004 | 0.009 | 4.6 | 6.33E-06*** |
|  | top-down | SAMy | -0.009 | -0.012 | -0.007 | -6.591 | 8.76E-11*** |
|  | top-down | audio env | 0.003 | 0 | 0.006 | 2.141 | 3.23E-02* |
|  | bottom-up | SAMy | -0.016 | -0.019 | -0.013 | -11.191 | 0.00E+00*** |
|  | bottom-up | audio env | -0.004 | -0.006 | -0.001 | -2.459 | 1.67E-02* |
|  | SAMy | audio env | 0.012 | 0.01 | 0.015 | 8.731 | 0.00E+00*** |
| Prefrontal | top-down | bottom-up | 0.012 | 0.007 | 0.017 | 4.508 | 5.62E-05*** |
|  | top-down | SAMy | 0.002 | -0.003 | 0.007 | 0.807 | 6.13E-01 |
|  | top-down | audio env | 0.01 | 0.005 | 0.016 | 3.984 | 5.07E-04*** |
|  | bottom-up | SAMy | -0.01 | -0.015 | -0.005 | -3.7 | 1.29E-03** |
|  | bottom-up | audio env | -0.001 | -0.006 | 0.004 | -0.523 | 7.67E-01 |
|  | SAMy | audio env | 0.008 | 0.003 | 0.013 | 3.177 | 8.11E-03** |
| Cingulate | top-down | bottom-up | 0.012 | 0.002 | 0.023 | 2.285 | 7.44E-02 |
|  | top-down | SAMy | 0.005 | -0.005 | 0.016 | 0.954 | 5.67E-01 |
|  | top-down | audio env | 0.012 | 0.001 | 0.023 | 2.229 | 7.75E-02 |
|  | bottom-up | SAMy | -0.007 | -0.018 | 0.003 | -1.331 | 3.54E-01 |
|  | bottom-up | audio env | 3.04E-04 | -0.011 | 0.01 | -0.056 | 9.55E-01 |
|  | SAMy | audio env | 0.007 | -0.004 | 0.017 | 1.275 | 3.79E-01 |
| Orbitofrontal | top-down | bottom-up | 0.01 | -0.001 | 0.02 | 1.847 | 1.69E-01 |
|  | top-down | SAMy | -0.002 | -0.012 | 0.008 | -0.398 | 7.92E-01 |
|  | top-down | audio env | 0.006 | -0.004 | 0.017 | 1.226 | 4.00E-01 |
|  | bottom-up | SAMy | -0.012 | -0.022 | -0.002 | -2.245 | 7.75E-02 |
|  | bottom-up | audio env | -0.003 | -0.014 | 0.007 | -0.62 | 7.14E-01 |
|  | SAMy | audio env | 0.009 | -0.002 | 0.019 | 1.625 | 2.40E-01 |
| Paracentral | top-down | bottom-up | 0.013 | 1.40E-04 | 0.026 | 1.939 | 1.43E-01 |
|  | top-down | SAMy | 0.001 | -0.012 | 0.014 | 0.175 | 9.19E-01 |
|  | top-down | audio env | 0.006 | -0.007 | 0.019 | 0.911 | 5.88E-01 |
|  | bottom-up | SAMy | -0.012 | -0.025 | 0.001 | -1.763 | 1.91E-01 |
|  | bottom-up | audio env | -0.007 | -0.02 | 0.006 | -1.028 | 5.21E-01 |
|  | SAMy | audio env | 0.005 | -0.008 | 0.018 | 0.735 | 6.30E-01 |
| Parietal | top-down | bottom-up | 0.004 | -0.006 | 0.015 | 0.776 | 6.13E-01 |
|  | top-down | SAMy | -0.027 | -0.037 | -0.016 | -4.866 | 1.37E-05*** |
|  | top-down | audio env | -0.001 | -0.011 | 0.01 | -0.102 | 9.50E-01 |
|  | bottom-up | SAMy | -0.031 | -0.042 | -0.02 | -5.643 | 2.51E-07*** |
|  | bottom-up | audio env | -0.005 | -0.016 | 0.006 | -0.879 | 5.99E-01 |
|  | SAMy | audio env | 0.026 | 0.015 | 0.037 | 4.764 | 1.90E-05*** |
| MedialWall & Subcortex | top-down | bottom-up | 0.001 | -0.007 | 0.009 | 0.273 | 8.56E-01 |
|  | top-down | SAMy | -0.007 | -0.015 | 0.001 | -1.753 | 1.91E-01 |
|  | top-down | audio env | -0.001 | -0.009 | 0.007 | -0.16 | 9.19E-01 |
|  | bottom-up | SAMy | -0.008 | -0.016 | 2.75E-04 | -2.027 | 1.22E-01 |
|  | bottom-up | audio env | -0.002 | -0.01 | 0.006 | -0.433 | 7.87E-01 |
|  | SAMy | audio env | 0.007 | -0.002 | 0.015 | 1.594 | 2.47E-01 |
| Superior Temporal-Insula | top-down | bottom-up | -0.002 | -0.009 | 0.005 | -0.449 | 7.87E-01 |
|  | top-down | SAMy | -0.041 | -0.048 | -0.034 | -11.444 | 0.00E+00*** |
|  | top-down | audio env | -0.011 | -0.018 | -0.004 | -2.971 | 1.41E-02* |
|  | bottom-up | SAMy | -0.04 | -0.047 | -0.033 | -10.995 | 0.00E+00*** |
|  | bottom-up | audio env | -0.009 | -0.016 | -0.002 | -2.522 | 4.67E-02* |
|  | SAMy | audio env | 0.031 | 0.023 | 0.038 | 8.473 | 0.00E+00*** |
| Lateral Temporal | top-down | bottom-up | 0.005 | -0.002 | 0.012 | 1.469 | 2.94E-01 |
|  | top-down | SAMy | -0.009 | -0.016 | -0.002 | -2.389 | 5.96E-02 |
|  | top-down | audio env | 0.001 | -0.006 | 0.008 | 0.386 | 7.92E-01 |
|  | bottom-up | SAMy | -0.014 | -0.021 | -0.007 | -3.857 | 7.64E-04*** |
|  | bottom-up | audio env | -0.004 | -0.011 | 0.003 | -1.083 | 4.92E-01 |
|  | SAMy | audio env | 0.01 | 0.003 | 0.017 | 2.775 | 2.37E-02* |
| Ventral Temporal | top-down | bottom-up | 0.006 | -0.002 | 0.015 | 1.418 | 3.13E-01 |
|  | top-down | SAMy | -0.007 | -0.016 | 0.002 | -1.544 | 2.63E-01 |
|  | top-down | audio env | 0.004 | -0.005 | 0.013 | 0.857 | 6.02E-01 |
|  | bottom-up | SAMy | -0.014 | -0.023 | -0.005 | -2.961 | 1.41E-02* |
|  | bottom-up | audio env | -0.003 | -0.012 | 0.006 | -0.561 | 7.50E-01 |
|  | SAMy | audio env | 0.011 | 0.002 | 0.02 | 2.401 | 5.96E-02 |
| Occipital | top-down | bottom-up | -0.001 | -0.019 | 0.018 | -0.06 | 9.55E-01 |
|  | top-down | SAMy | -0.008 | -0.026 | 0.01 | -0.834 | 6.07E-01 |
|  | top-down | audio env | -0.005 | -0.023 | 0.014 | -0.488 | 7.82E-01 |
|  | bottom-up | SAMy | -0.007 | -0.025 | 0.011 | -0.773 | 6.13E-01 |
|  | bottom-up | audio env | -0.004 | -0.022 | 0.014 | -0.428 | 7.87E-01 |
|  | SAMy | audio env | 0.003 | -0.015 | 0.021 | 0.345 | 8.11E-01 |

**Supp. Table 2.** Top-down and bottom-up simulation parameters.

| Simulation Parameters |  |  |  |
| --- | --- | --- | --- |
| Parameter | Variable | Value | Description |
| Top-Down Dynamic Attention Model |  |  |  |
| tone onset | c | trial-dependent | time of expected tone onset, center of gaussian |
| tone number | n | trial-dependent | number of tones in the sequence |
| inter-onset interval (IOI) | b | trial-dependent | ms, time between expected isochronous tone onsets |
| gaussian width | w | 100 | ms, gaussian of temporal excitability centered around tone onset |
| gaussian height | a | 1 | maximum firing rate |
| learning rate | l | 0.15 | narrows gaussian with each successive tone |
| sensitization rate | s | 0.2 | heightens gaussian with each successive tone |
| Bottom-Up Auditory Nerve Response Model |  |  |  |
| response amplitude | $a_r$ | 1 | maximum stimulus-evoked firing rate |
| response decay | $\tau_r$ | 10 | time constant of evoked response |
| short term amplitude | $a_{st}$ | 0.25 | post-evoked short term firing rate |
| short term decay | $\tau_{st}$ | 100 | time constant of short term response |
| plateau amplitude | $a_{base}$ | 0.5 | steady state firing probability |

